# A genome-wide catalog of sites where peripheral DNA methylation reflects the brain

**DOI:** 10.64898/2026.10.23.761276

**Authors:** Shota Nishitani, Shizhong Han, Takaya Ishii, Akiyoshi Shimura, Nathan J. Phuong, Asahi Kato, Meagan Chambers, Tao Wang, Makoto Isozaki, Yoshifumi Higashino, Takahiro Yamauchi, Satoshi Kawajiri, Hidetaka Arishima, Toshiaki Kodera, Hirochika Imoto, Sadahiro Nomura, Kyosuke Yamanishi, Bun Aoyama, Ilgin Genc, Rika Sakuma, Hieu D. Nguyen, Nipun Gorantla, Elnaz Ghasemi, Akiko Yao, Sawa Kurata, Takashi X. Fujisawa, Patricia R. Braun, Takaaki Nagao, Takehiko Yamanashi, Akemi Tomoda, Kenichiro Kikuta, Tadao Ooka, Gerald A. Grant, Jody E. Hooper, Matthew A. Howard, Hiroto Kawasaki, James B. Potash, Michael P. Snyder, Gen Shinozaki

**Author notes:** These authors are co-corresponding authors. Address correspondence to: Gen Shinozaki, Division of Geriatrics, Department of Family Medicine, School of Medicine, University of California, Irvine, Irvine, CA 92617, USA;., Shota Nishitani, Division of Geriatrics, Department of Family Medicine, School of Medicine, University of California, Irvine, Irvine, CA 92617, USA;., Michael P. Snyder, Department of Genetics, School of Medicine, Stanford University, Palo Alto, CA 94304, USA;.

## Abstract

Epigenetic studies of psychiatric and neurological disorders rely largely on peripheral tissues, yet where peripheral DNA methylation (DNAm) reflects the brain remains unclear. We generated a paired brain and peripheral DNAm dataset from 148 donors in three cohorts and combined it with five published cohorts for consensus brain–peripheral correlations. By cross-validation across cohorts, we defined brain epigenome shared sites (BrESS), more than 100,000 CpGs per tissue at which peripheral DNAm reproducibly predicts brain DNAm. BrESS lay at promoters active in both brain and periphery and in housekeeping genes, and most were not explained by genetic effects. BrESS did not contribute to the heritability of seven disorders, and CpGs from epigenome-wide association studies of these disorders were not enriched at BrESS. Peripheral DNAm reflects the brain mainly in a regulatory compartment shared across tissues, outside which disease signals largely lie, and BrESS separate peripheral signals attributable to the brain from others.

## Introduction

Epigenetic mechanisms, particularly DNA methylation (DNAm), are central to gene regulation in the human body including the brain and are implicated in the pathophysiology of psychiatric and neurological disorders^1–3^. DNAm in post-mortem brain has been examined in schizophrenia^4^, major depressive disorder^5,6^, autism spectrum disorder^7^, Alzheimer’s disease^8,9^ and Parkinson’s disease^10^, but such studies can be performed only under limited conditions and in limited numbers of donors. Because DNAm is highly tissue-specific^11,12^ and the organ of interest in these disorders is the brain, understanding their epigenetic basis ideally requires brain tissue. Brain tissue, however, is largely inaccessible in living humans, and this has been a major barrier to progress in the epigenetics of these disorders. As a result, the great majority of human epigenome-wide association studies (EWAS) in this field rely on peripheral tissues, mainly blood, largely on the untested assumption that peripheral DNAm at least partially reflects the brain epigenome^13^.

Whether a CpG identified in a peripheral EWAS of a disorder can inform epigenetic mechanisms in the brain, or can be interpreted only as a peripheral mechanism or biomarker unrelated to the brain, therefore remains unknown.

To address this uncertainty, correlation resources using paired brain and peripheral samples from the same individuals have been developed. Following early comparisons of blood with post-mortem brain^14,15^, IMAGE-CpG^16^ extended this approach to brain tissue from living individuals and to three peripheral tissues (blood, buccal epithelium and saliva). These resources, however, have remained limited in three respects. (i) Individual cohorts are small. (ii) Results have not been integrated across cohorts. (iii) None of these resources has defined, genome-wide and with a validated criterion, the set of CpG sites at which peripheral DNAm reliably reflects the brain.

Here, we addressed these limitations. (i) We generated paired DNAm data from 148 donors across three newly assembled cohorts spanning adult biopsy, adult autopsy and pediatric samples, covering multiple brain regions and three peripheral tissues (IMAGE-CpG2). This dataset is approximately seven times larger than the original IMAGE-CpG cohort and the largest of its kind to date. (ii) We combined these data with five previously published cohorts^14–18^ to provide consensus estimates of brain–peripheral correlation (COMPASS). (iii) Using the new cohorts, we defined brain epigenome shared sites (BrESS), more than 100,000 CpGs per peripheral tissue at which peripheral DNAm reproducibly reflects brain DNAm in cross-validation across cohorts (TRACE-CpG). Having BrESS as a validated set allowed us, for the first time, to characterize these sites comprehensively, including their genomic and chromatin context, the genes they belong to and their genetic control, and to examine their relationship to the heritability of psychiatric and neurological disorders and to CpGs reported in published EWAS. Together, these resources provide a reference framework for interpreting peripheral DNAm studies of the human brain.

## Results

### A multi-cohort resource of 148 paired brain–peripheral DNAm profiles

To determine where and how strongly peripheral DNAm reflects the brain, we built three resources (Fig. 1). First, a brain–peripheral DNAm correlation atlas based on newly generated paired data from 148 donors in three cohorts (adult surgical biopsy, DB6; adult autopsy, DB7; pediatric, DB8), covering four brain regions and three peripheral tissues (blood, buccal epithelium and saliva) (IMAGE-CpG2; Irvine Methylation Array Graphing Experiment for Comparison of peripheral tissue and Gray matter, version 2; donor characteristics in Table 1). Second, a meta-analytic platform that integrates IMAGE-CpG2 with five previously published cohorts to provide consensus correlation estimates (COMPASS; COrrelation Methylation Platform for Association Search and Synthesis). Third, a linear mixed-effects modeling framework built on IMAGE-CpG2 that defines brain epigenome shared sites (BrESS) (TRACE-CpG; Trans-tissue Regression Approximation of Central Epigenome from Correlated peripheral and Gray-matter).

**Figure 1.**
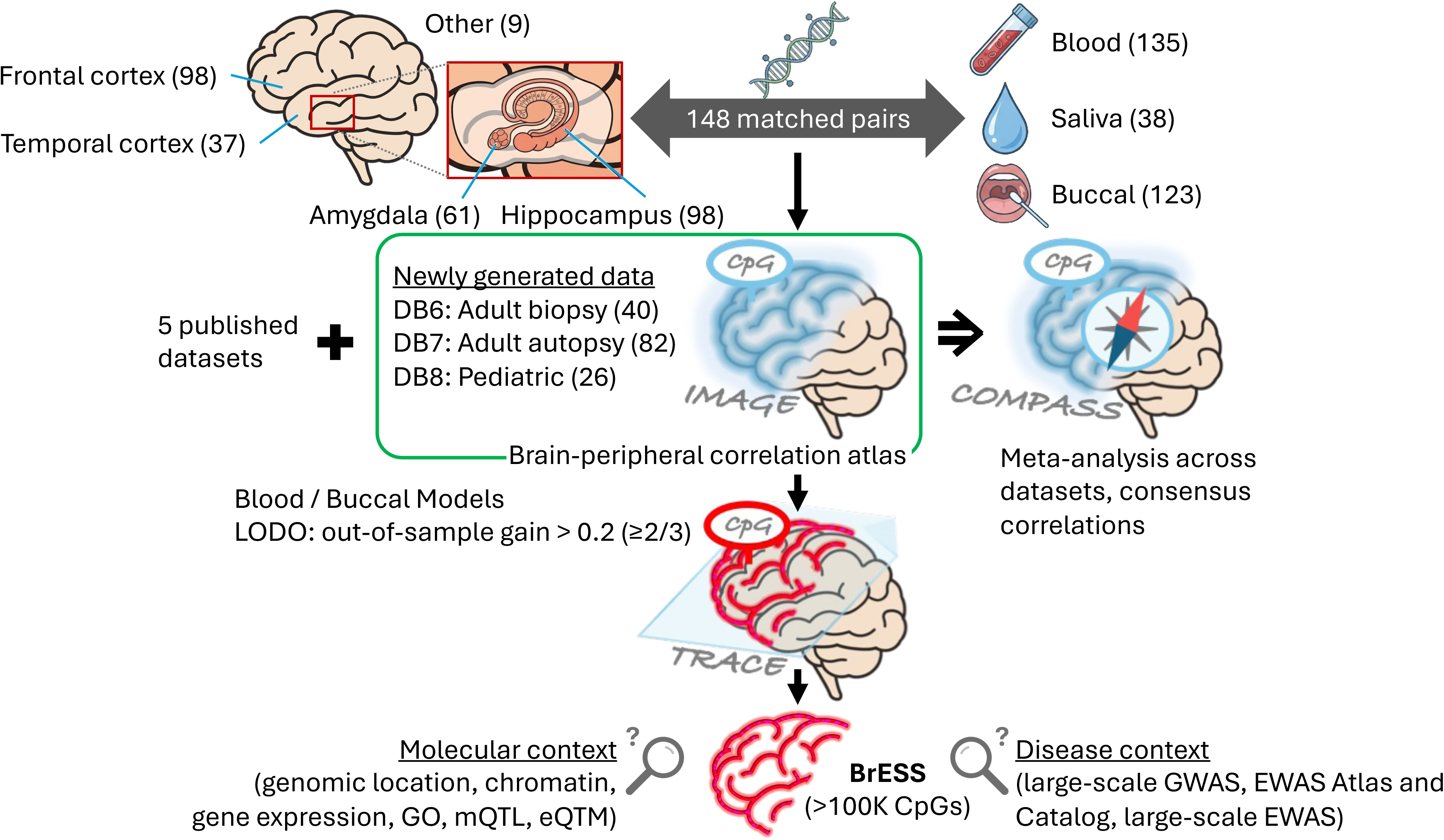
Overview of the study design and analytical framework. A total of 148 subjects with matched brain and peripheral tissue samples were assembled across three newly generated datasets: DB6 (adult biopsy, n = 40), DB7 (adult autopsy, n = 82) and DB8 (pediatric, n = 26). Brain tissues included frontal cortex (98 samples), temporal cortex (37), amygdala (61), hippocampus (98) and other regions (9). Peripheral tissues included blood (135), saliva (38) and buccal epithelium (123). These paired data were used to construct IMAGE-CpG2, a brain–peripheral DNAm correlation atlas. Integration with five previously published datasets (DB1–5) by random-effects meta-analysis yielded COMPASS, a resource of consensus brain–peripheral correlations. Using TRACE-CpG, a cross-validated prediction framework built on IMAGE-CpG2, we identified CpG sites at which peripheral DNAm reproducibly captures brain DNAm, termed brain epigenome shared sites (BrESS), from blood and buccal epithelium. BrESS were then characterized in their molecular and disease contexts.

**Table 1.** Summary of previously published and newly generated cross-tissue DNAm cohorts.

|  | DB1<br>Hannon<br>ND brain<br>bank | DB2<br>Edgar<br>Sudden<br>death<sup>a</sup> | DB3<sup>b</sup><br>Braun<br>Epilepsy<br>surgery | DB4<br>Sommerer<br>AD research<br>center | DB5<sup>c</sup><br>Nishitani<br>Brain surgery<sup>d</sup> | DB6<br>(biopsy)<br>Epilepsy | DB7<br>(autopsy)<br>Various<br>causes | DB8<br>(pediatric)<br>Epilepsy | IMAGE-CpG2<br>Integrated |
| --- | --- | --- | --- | --- | --- | --- | --- | --- | --- |
| Status | Published | Published | Published | Published | Updated | New | New | New | New |
| *n* | 71–75 | 15 | 6 | 108 | 35 | 40 | 82 | 26 | 148 |
| Array |  |  |  |  |  |  |  |  |  |
| 450K | 71–75 | 15 | 6 | 0 | 0 | 0 | 0 | 0 | 0 |
| EPICv1/v2/v2C | 0/0/0 | 0/0/0 | 0/0/0 | 108/0/0 | 35/0/0 | 40/0/0 | 26/19/37 | 23/0/3 | 89/19/40 |
| Bio/Auto-psy | 0/71–75 | 0/15 | 6/0 | 0/108 | 35/0 | 40/0 | 0/82 | 22/4 | 62/86 |
| Ancestry |  |  |  |  |  |  |  |  |  |
| European | 71–75 | 15 | 6 | 108 | 0 | 37 | 71 | 13 | 121 |
| East Asian | 0 | 0 | 0 | 0 | 35 | 0 | 4 | 5 | 9 |
| Other | 0 | 0 | 0 | 0 | 0 | 3 | 7 | 8 | 18 |
| Age (y) | 71–105<sup>e</sup> | 15–87<sup>f</sup> | 37 (18–60) | 72 | 60 (11–82) | 37 (19–64) | 66 (21–97) | 8 (0.3–18) | 48 (0.3–97) |
| Sex (M/F) | 26/49 | 13/4<sup>f</sup> | 5/1 | NA<sup>g</sup> | 16/19 | 22/18 | 48/34 | 15/11 | 85/63 |
| Tissues |  |  |  |  |  |  |  |  |  |
| PFC/BA10 | 74 | 15 | 3 | 101 | 15 | 5 | 81 | 12 | 98 |
| TC/STG/BA20 | 75 | 15 | 1 | 0 | 16 | 27 | 0 | 10 | 37 |
| AMY | 0 | 0 | 1 | 0 | 0 | 14 | 45 | 2 | 61 |
| HIPP | 0 | 0 | 2 | 0 | 1 | 19 | 77 | 2 | 98 |
| Other | 142 | 15 | 1 | 0 | 3 | 5 | 0 | 4 | 9 |
| BL/EP/SA | 75/0/0 | 15/0/0 | 6/0/6 | 0/101/0 | 35/35/35 | 40/32/32 | 72/81/0 | 23/10/6 | 135/123/38 |
| Total samples | 366 | 60 | 20 | 202 | 140 | 174 | 356 | 69 | 599 |

### Brain–peripheral DNAm correlations are reproducible across biopsy, autopsy and pediatric cohorts

Multidimensional scaling (MDS) of genome-wide DNAm profiles showed tissue identity as the dominant axis of variation in each of the three cohorts and in the integrated dataset, with brain samples separated from the three partially overlapping peripheral tissues (Fig. 2A). Within each tissue, biopsy and autopsy samples did not separate, indicating that DNAm profiles were comparable across collection methods after preprocessing. Because IMAGE-CpG2 includes the hippocampus and amygdala, we examined whether brain–peripheral correlations differed by brain region. In DB7, where multiple regions were available within individuals, the distributions of CpG-wise correlations were similar across prefrontal cortex, hippocampus and amygdala (Supplementary Fig. 1A). Correlations at individual CpGs were highly concordant between any two regions (Pearson r = 0.71–0.79 for blood and buccal epithelium; Supplementary Fig. 1B), although the regions themselves were only moderately correlated (Supplementary Table 1). Regional DNAm differences thus have limited impact on the relationship with the periphery, and we used pan-brain DNAm, the within-subject average across regions, as the primary measure of brain DNAm.

**Figure 2.**
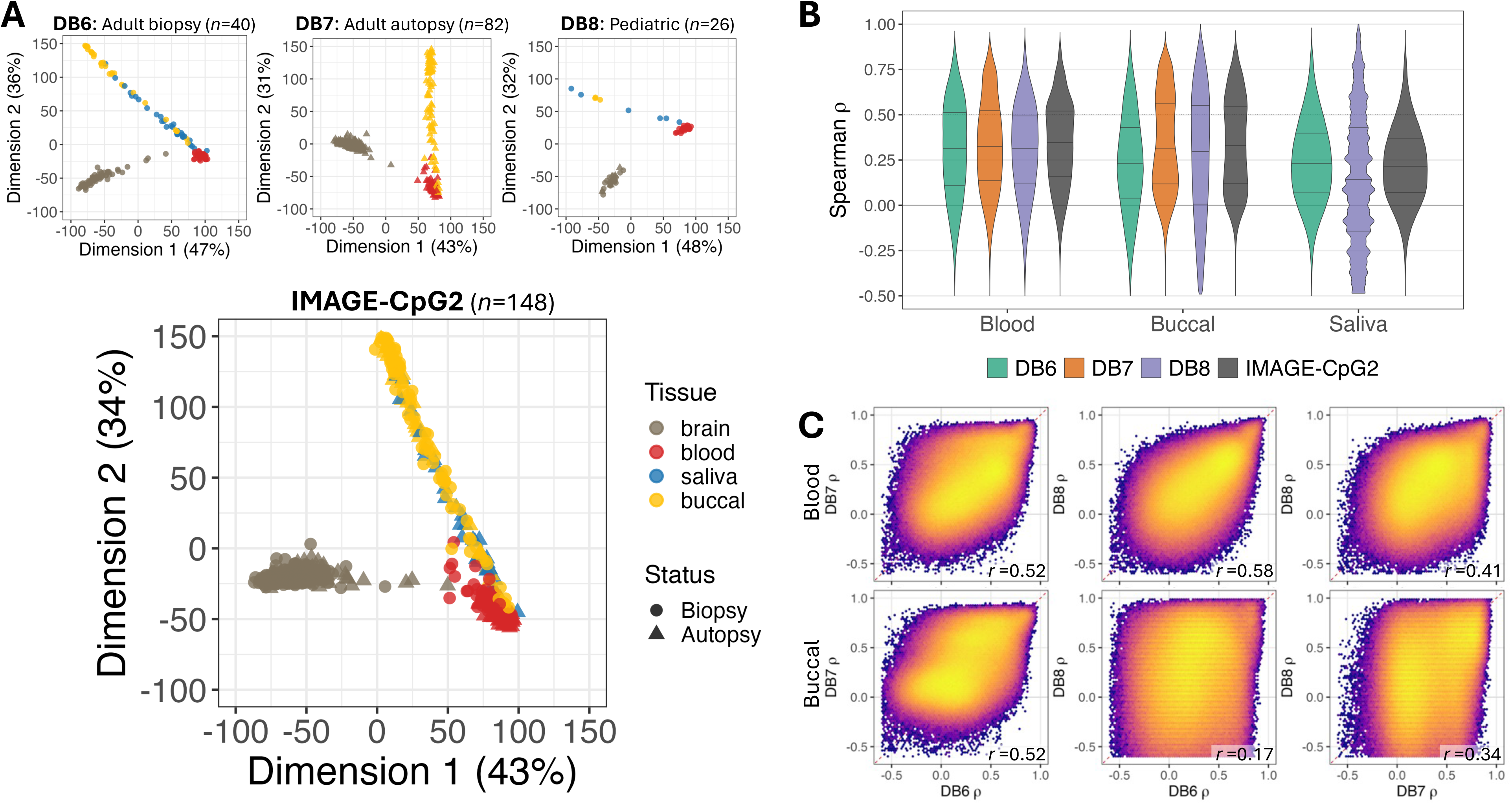
Characterization of the IMAGE-CpG2 resource across collection methods and age groups. **(A)** Multidimensional scaling (MDS) plots of genome-wide DNAm profiles for DB6 (Adult-Biopsy, *n* = 40), DB7 (Adult-Autopsy, *n* = 82), DB8 (Pediatric, *n* = 26), and the integrated IMAGE-CpG2 dataset (*n* = 148). Each point represents a sample, colored by tissue (brain, blood, saliva, buccal epithelium) and shaped by collection method (biopsy, autopsy). Percentage of variance explained by each dimension is shown in parentheses. **(B)** Violin plots of CpG-wise Spearman correlations (ρ) between pan-brain and peripheral tissue DNAm (blood, buccal epithelium, saliva) for each dataset (DB6, DB7, DB8) and the integrated IMAGE-CpG2 cohort. Pan-brain DNAm was computed by averaging across all available brain regions within each subject. Horizontal lines within violins indicate the 25th, 50th, and 75th percentiles. Solid horizontal line at ρ = 0 and dotted line at ρ = 0.5 serve as references. DB7 did not include saliva samples. Summary statistics are in Supplementary Table 2. **(C)** Pairwise concordance of CpG-wise ρ values between datasets for blood (top row) and buccal epithelium (bottom row). Each panel compares ρ values computed in one dataset (x-axis) versus another (y-axis) across 687,873 CpG sites. Red dashed line indicates *y* = *x*. Pearson *r* in the bottom-right corner quantifies the agreement of ρ values between datasets. Saliva comparisons are not shown as DB7 lacked saliva samples.

For each CpG site, we then computed Spearman’s correlation (ρ) between pan-brain DNAm and peripheral tissue DNAm across paired subjects. The resulting distributions of CpG-wise ρ were broadly similar across DB6 (adult biopsy), DB7 (adult autopsy) and DB8 (pediatric) for blood and buccal epithelium. In the integrated IMAGE-CpG2 dataset the majority of CpG sites showed positive brain–peripheral correlation, with median ρ of 0.35, 0.33 and 0.22 for blood, buccal epithelium and saliva, respectively. The fraction of sites with ρ > 0.2 was 70% for blood, 65% for buccal epithelium and 53% for saliva, while a smaller fraction exhibited strong positive brain–peripheral correlations (ρ > 0.5: 10–31%) (Fig. 2B; Supplementary Table 2). Saliva was not collected in DB7 and was limited in DB8 (n = 6), precluding robust cohort-level estimates.

We next assessed whether site-level correlations were reproducible across cohorts. Pairwise comparisons of CpG-wise ρ showed moderate concordance for blood across all cohort pairs (Pearson r = 0.41–0.58), whereas buccal epithelium showed more variable agreement, lowest between DB6 and DB8 (Fig. 2C). Comparing biopsy and autopsy directly (Δρ = ρ_DB6 − ρ_DB7), the distribution for blood was tightly centered near zero, whereas for buccal epithelium it showed a small systematic shift away from zero rather than increased noise (Supplementary Fig. 1C,D). Thus, blood correlations differed little between biopsy and autopsy, whereas buccal correlations showed a small systematic change. The broad pattern of brain–peripheral correlation was nonetheless reproducible across the adult and pediatric cohorts, supporting their integration as a single dataset (IMAGE-CpG2).

### A meta-analytic consensus atlas of brain–peripheral DNAm correlations across multiple cohorts

To integrate evidence across cohorts, we developed COMPASS, which pools the per-cohort correlations at each CpG and estimates between-cohort heterogeneity (I²). In the brain–blood meta-analysis, shown here as a representative example, seven cohorts contributed (DB1–3 and DB5–8; Table 1; DB4 contributes only to the buccal meta-analysis) and 350,274 CpGs were analyzed. Heterogeneity varied substantially across CpGs (Fig. 3A). For 19.2% of CpGs it was very low (I² < 5%), indicating that the brain–blood correlation was consistent across cohorts despite differences in population, brain region, collection method and array platform, whereas 29.0% showed high heterogeneity (I² > 75%). Pooled correlations were predominantly positive (median ρ = 0.206, IQR 0.111–0.304; Fig. 3B) and tended to be stronger, not weaker, at higher-heterogeneity sites (Fig. 3C). Because a strong pooled correlation does not guarantee consistency across cohorts, COMPASS provides both measures for each CpG so that users can evaluate them jointly.

**Figure 3.**
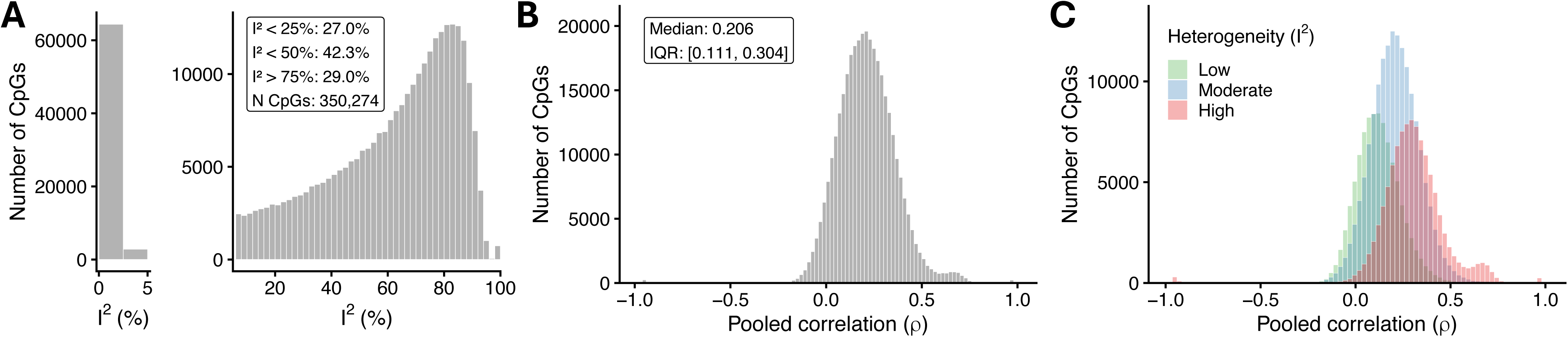
Meta-analytic integration of cross-tissue DNAm correlations across seven databases (COMPASS). Spearman’s correlations (ρ) between brain (pan-brain) and blood DNAm were computed for each CpG site within each database and integrated using a random-effects meta-analysis (REML). 350,274 CpG sites common to the contributing databases (DB1–3, DB5–8) were included. **(A)** Distribution of heterogeneity (I²) across CpG sites. The left sub-panel shows sites with I² < 5%; the right sub-panel shows the remaining sites. **(B)** Distribution of pooled correlation coefficients. **(C)** Distribution of pooled ρ stratified by heterogeneity: low (I² < 25%), moderate (25–75%), and high (> 75%).

### Genome-wide identification of brain epigenome shared sites (BrESS) by cross-validation

IMAGE-CpG2 and COMPASS describe brain–peripheral correlations one CpG at a time. To define the sites at which peripheral DNAm robustly reflects brain DNAm, we developed TRACE-CpG, which models brain DNAm from peripheral DNAm at each CpG, and used it to evaluate all 687,873 CpGs by leave-one-dataset-out cross-validation (LODO-CV) across the three IMAGE-CpG2 cohorts, measuring the out-of-sample gain in variance explained in brain DNAm by peripheral DNAm (R²gain). We defined BrESS as CpGs with R²gain > 0.2 in at least two of the three cycles (Fig. 4A), yielding 121,167 sites for blood and 109,385 for buccal epithelium, of which 76,208 were shared (Fig. 4B). Although R²gain was small at most CpGs, in each cycle 15–25% of CpGs for blood and 11–22% for buccal epithelium exceeded 0.2 (Fig. 4C; Supplementary Table 3). Saliva was not included because the available data did not permit LODO-CV (per-site results in Supplementary Data 1).

**Figure 4.**
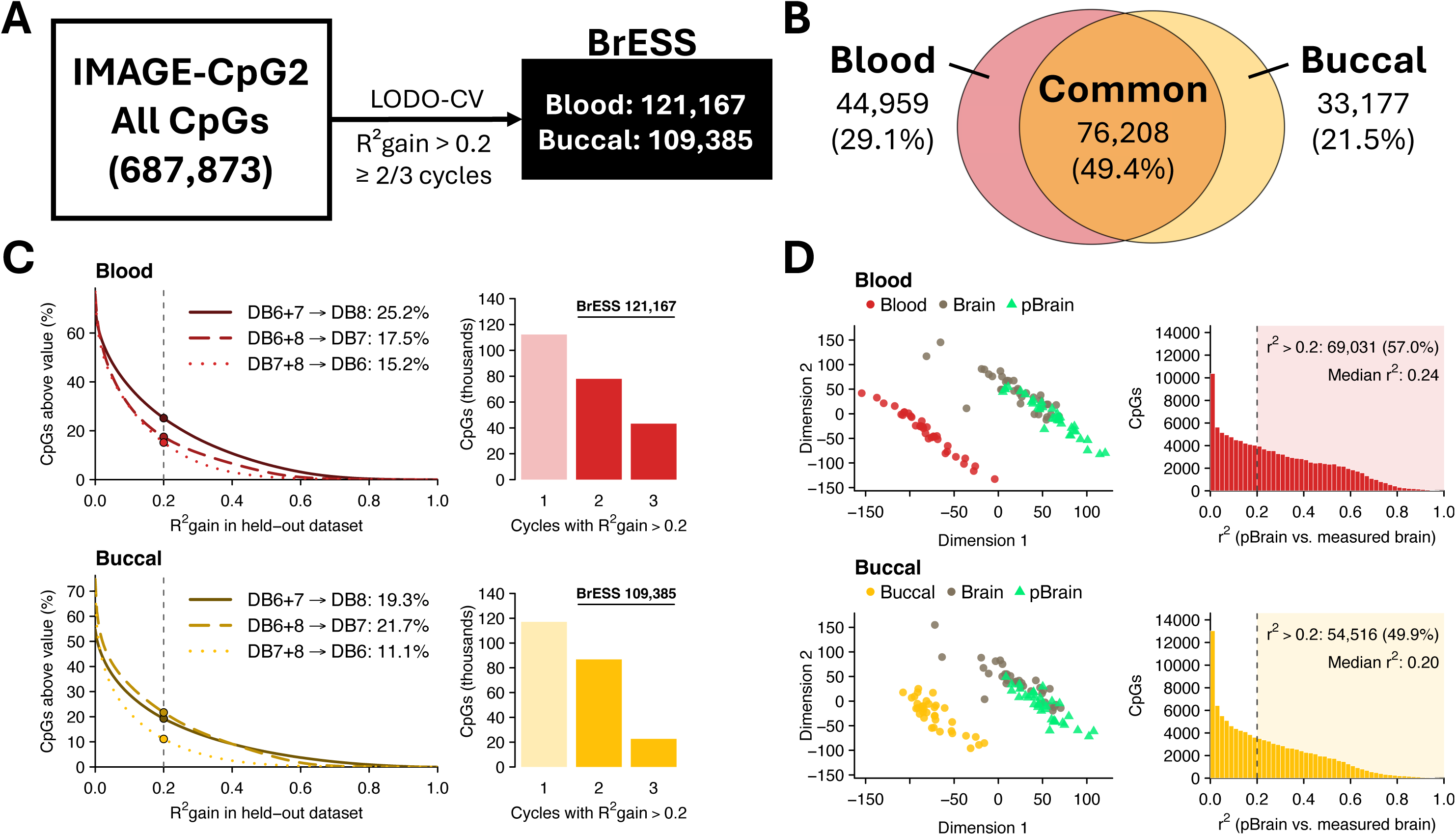
Identification and external validation of brain epigenome shared sites (BrESS). **(A)** Identification of BrESS in IMAGE-CpG2 by leave-one-dataset-out cross-validation (LODO-CV) across DB6–8. R²gain, gain in out-of-sample R² for brain DNAm from adding peripheral DNAm to a model with age and sex. **(B)** Overlap between blood and buccal BrESS. Percentages are relative to CpGs that are BrESS in either tissue. **(C)** R²gain in each LODO-CV cycle for blood (top) and buccal epithelium (bottom). Left: percentage of CpGs whose R²gain exceeds a given value; lines indicate cycles (training datasets → held-out dataset), and points mark the threshold of 0.2 (dashed line). Right: number of CpGs with R²gain > 0.2 in one, two or three cycles. **(D)** External validation in AMAZE-CpG2 (n = 35). Left: multidimensional scaling of measured peripheral DNAm, measured brain DNAm and predicted brain DNAm (pBrain) at BrESS. Right: distribution of r² between pBrain and measured brain DNAm at BrESS; shaded areas and numbers indicate CpGs with r² > 0.2. LODO-CV results are in Supplementary Table 3, and model fit and external validation are in Supplementary Tables 4 and 7.

### External validation of prediction at BrESS in an independent cohort

Although BrESS were selected by out-of-sample performance, the prediction coefficients were estimated on the full IMAGE-CpG2 data (Supplementary Data 2), so agreement within these data reflects in-sample fit (median variance in brain DNAm explained by peripheral DNAm at BrESS, 33% for blood and 43% for buccal epithelium; Supplementary Table 4). We therefore tested the coefficients in AMAZE-CpG2 (DB5; n = 35), an independent dataset not used at any stage of BrESS derivation or model fitting, which expands the previously published AMAZE-CpG^18^ from 19 to 35 subjects (Supplementary Note 1; Supplementary Tables 5 and 6; Supplementary Figs. 2 and 3). In MDS, predicted brain DNAm (pBrain) separated from the originating peripheral tissue and clustered with measured brain samples (Fig. 4D). The median r² between pBrain and measured brain DNAm was 0.24 for blood and 0.20 for buccal epithelium, with 57.0% and 49.9% of BrESS, respectively, exceeding 0.2 (Fig. 4D; Supplementary Table 4).

Based on this validation, we provide TRACE-CpG as a web tool and as code that compute pBrain at BrESS from user-supplied peripheral DNAm, age and sex (Supplementary Data 2). Because pBrain is a linear function of peripheral DNAm, it adds no information over the peripheral data in group-level comparisons. Rather, it indicates the level that brain DNAm is expected to take after its compressed variance is rescaled (Supplementary Table 7; Supplementary Fig. 4; Supplementary Note 2).

### BrESS are enriched at promoters active across tissues and in broadly expressed genes

To characterize what distinguishes BrESS, we compared BrESS with non-BrESS sites in their genomic and chromatin context and in the expression and function of the genes they belong to. Blood BrESS were preferentially located within CpG islands (43.3% vs 14.2% of non-BrESS, 3.0-fold) and near transcription start sites (TSS200, 2.9-fold; first exon, 2.5-fold), with a closely matching profile for buccal BrESS, whereas non-BrESS sites were more common in open-sea and intergenic regions and gene bodies (Fig. 5A; Supplementary Table 8).

**Figure 5.**
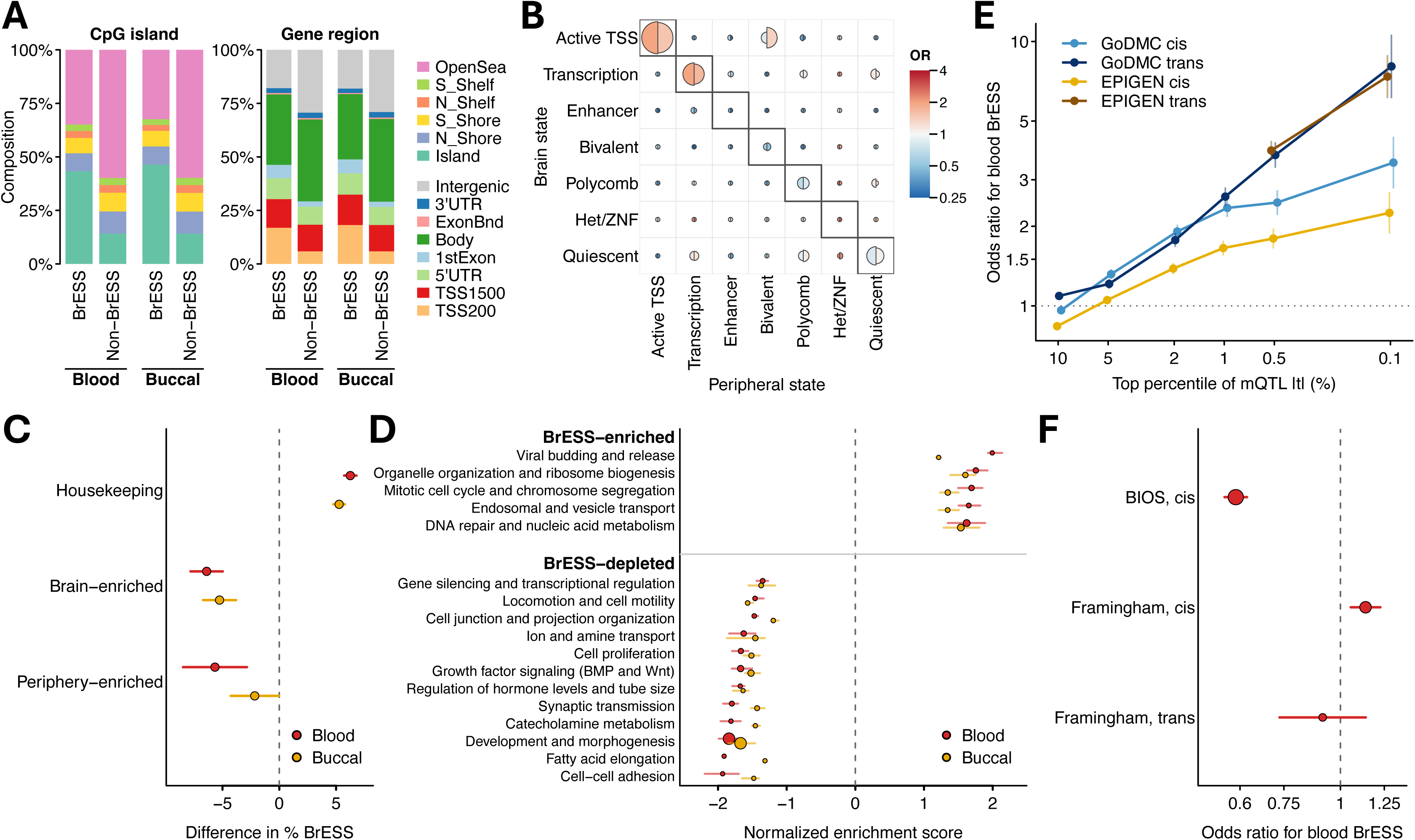
Genomic and functional features of BrESS. **(A)** Distribution of BrESS and non-BrESS CpGs by relation to CpG islands (left) and gene region (right) in blood and buccal samples. **(B)** Chromatin states of BrESS in brain and peripheral tissues, based on the Roadmap Epigenomics Project. Rows give the chromatin state in brain and columns give the state in the matched peripheral reference (blood or buccal epithelium). Boxed cells on the diagonal mark CpGs in the same state in both tissues. In each cell, the left half-circle shows blood BrESS and the right half-circle shows buccal BrESS. Color indicates the odds ratio for BrESS, and half-circle size reflects the percentage of BrESS in that cell. **(C)** Difference in the percentage of BrESS between genes in each expression class, defined from GTEx, and all other genes. Periphery-enriched genes are blood-enriched genes for blood BrESS and epithelium-enriched genes for buccal BrESS. Horizontal lines indicate 95% confidence intervals. **(D)** Gene Ontology biological process clusters. Each row is a cluster of terms defined in blood BrESS. Circles show the mean normalized enrichment score of the terms in each cluster, horizontal lines show the interquartile range, and circle size reflects the number of terms. Positive scores indicate functions whose genes carry more BrESS. **(E)** Odds ratios for blood BrESS among CpGs ranked in the top percentiles of methylation QTL strength, shown separately for cis and trans effects in GoDMC and EPIGEN. Vertical lines indicate 95% confidence intervals. The y-axis is on a log scale. **(F)** Odds ratios for blood BrESS among CpGs with blood expression QTM associations from BIOS and the Framingham Heart Study. Horizontal lines indicate 95% confidence intervals, and circle size reflects the number of eQTM CpGs. The x-axis is on a log scale. In C and D, red indicates blood BrESS and yellow indicates buccal BrESS. Odds ratios in B, E and F and differences in C are adjusted for CpG island relation and gene region. Dashed lines mark no difference (0) or an odds ratio of 1. Numerical results are in Supplementary Tables 8 (A), 9 (B), 10 (C), 11 and 12 (D), 13 (E) and 14 (F), and GO clusters are shown in Supplementary Fig. 5.

We next examined chromatin states in Roadmap Epigenomics reference epigenomes^19^ of the brain and of a peripheral reference for each tissue (peripheral blood mononuclear cells for blood BrESS and keratinocytes for buccal BrESS). BrESS were enriched at CpGs in the same active state in the brain and the peripheral reference, namely active TSS (35.5% of blood BrESS vs 12.0% of non-BrESS) and transcribed regions, with odds ratios (ORs) of 1.5–2.0 after adjustment for genomic location, and were depleted at CpGs in enhancer, bivalent or Polycomb-repressed states (Fig. 5B; Supplementary Table 9).

Consistent with this, genes expressed in all tissues in GTEx^20^ carried more BrESS than other genes (+6.3 and +5.3 percentage points for blood and buccal BrESS), whereas brain-enriched genes (−6.4 and −5.3 points) and periphery-enriched genes (−5.7 and −2.2 points) carried fewer (Fig. 5C; Supplementary Table 10). Gene Ontology analysis showed the same contrast. Terms enriched in blood BrESS genes formed clusters of housekeeping processes, such as DNA repair and nucleic acid metabolism, organelle organization and the mitotic cell cycle, whereas depleted terms formed clusters such as development and morphogenesis (67% of depleted terms), synaptic transmission and growth factor signaling (BMP and Wnt) (Fig. 5D; Supplementary Fig. 5). Most of these terms showed the same direction for buccal BrESS (Fig. 5D; Supplementary Tables 11 and 12).

### Most BrESS are not explained by mQTL effects and are not enriched among eQTM CpGs

We further examined whether BrESS reflect genetically controlled DNAm, since brain–peripheral covariation could arise from shared genetic sequence. When CpGs were ranked by the strength of their mQTLs in blood, blood BrESS were enriched among the CpGs with the strongest effects (location-adjusted ORs in the top 0.1%: cis, 3.5 in GoDMC^21^ and 2.2 in EPIGEN^22^; trans, 8.1 and 7.4), but this enrichment diminished as the threshold was relaxed and had largely disappeared by the top 10% (Fig. 5E; Supplementary Table 13). These results indicate that brain–peripheral coupling attributable to strong mQTL effects is confined to a small set of extreme sites, and that the majority of BrESS instead capture coupling that is not accounted for by known mQTL effects. Consistent with this, CpGs within CoRSIVs, regions of systemic inter-individual variation under strong genetic influence^23^, were strongly enriched for both blood and buccal BrESS (location-adjusted OR 6.75 and 6.88) but accounted for less than 1% of BrESS.

We next asked whether BrESS coincide with CpGs whose DNAm is associated with gene expression in blood (eQTM CpGs). Blood BrESS were depleted among cis eQTM CpGs in BIOS^24^ (location-adjusted OR 0.59) and showed ORs of 1.14 for top cis and 0.91 for trans eQTM CpGs in the Framingham Heart Study^25^ (Fig. 5F; Supplementary Table 14). Thus, BrESS were not concentrated at eQTM CpGs, and the only OR above 1 (1.14) indicated at most weak enrichment.

### Contribution of BrESS to the heritability of psychiatric and neurological disorders

We then turned to psychiatric and neurological disorders. Using stratified LD score regression^26,27^ with GWAS of seven disorders^28–34^, we first asked whether SNPs near BrESS contribute to SNP heritability beyond baseline genomic annotations, which include promoter and transcription start site annotations, and all assessed CpGs (Fig. 6A; Supplementary Tables 15 and 16). The standardized effect of BrESS (τ*) did not differ significantly from zero for any disorder, for either blood or buccal BrESS (all P ≥ 0.17), although estimates for Parkinson’s disease and amyotrophic lateral sclerosis were imprecise owing to their low heritability (heritability z score < 7). Among educational attainment and ten non-brain traits, τ* was nominally negative for rheumatoid arthritis, inflammatory bowel disease and coronary artery disease with blood BrESS and for inflammatory bowel disease with buccal BrESS (P = 0.017–0.032), but none of these remained significant after correction for multiple testing (Fig. 6B; Supplementary Table 16).

**Figure 6.**
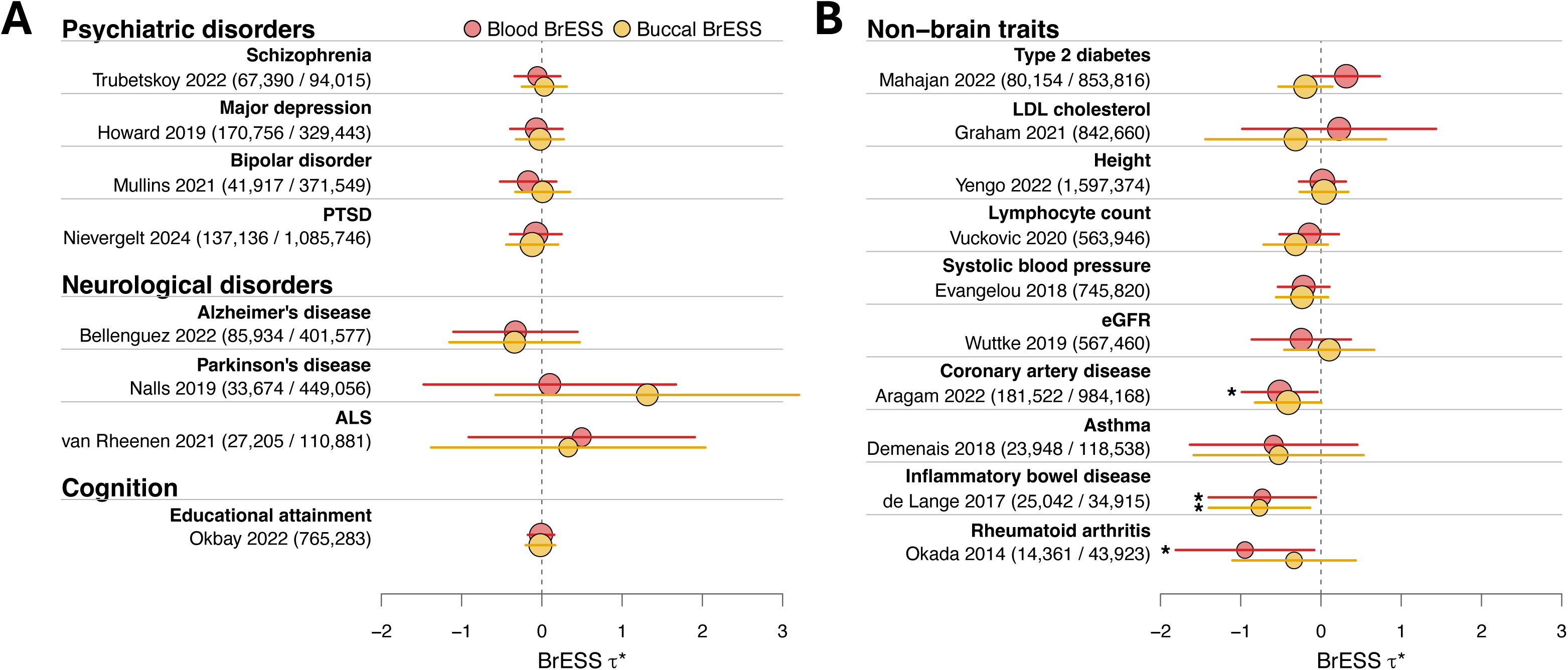
Contribution of BrESS to SNP heritability of psychiatric and neurological disorders and other traits. Contribution of BrESS to SNP heritability was estimated by stratified LD score regression. **(A)** Psychiatric disorders, neurological disorders and educational attainment. **(B)** Non-brain traits, ordered by τ* for blood BrESS. Each row shows one GWAS. Circles indicate the standardized effect of BrESS (τ*) beyond baseline genomic annotations and all assessed CpGs, and horizontal lines indicate 95% confidence intervals. Red circles show blood BrESS and yellow circles show buccal BrESS. Asterisks indicate nominal P < 0.05; none remained significant after Benjamini–Hochberg correction. The vertical dashed line marks a τ* of 0. Circle size reflects study sample size on a logarithmic scale, and numbers in parentheses give cases / controls or, for quantitative traits, the total sample size. GWAS sources and full results are in Supplementary Tables 15 and 16.

### Overlap of BrESS with CpGs reported in EWAS Atlas and EWAS Catalog studies

We next asked whether CpGs reported by published EWAS of the brain, and by peripheral EWAS of brain-related traits, are enriched for BrESS. For EWAS of brain, blood and saliva or buccal tissue in the EWAS Atlas^35,36^ and the EWAS Catalog^37^, we estimated for each study the OR for BrESS among the reported CpGs, adjusted for genomic location, and pooled the estimates by trait group (Fig. 7A and Supplementary Fig. 6; Supplementary Tables 17–19). Pooled ORs varied widely between trait groups. Among brain EWAS, CpGs reported for some traits were enriched for blood BrESS, including frontotemporal lobar degeneration and multiple system atrophy (pooled OR 4.45, 95% CI 3.38–5.84) and alcohol use disorder (1.88, 1.52–2.33), whereas those for others, such as multiple sclerosis (0.13, 0.07–0.23), were depleted (Fig. 7A; Supplementary Table 17). These estimates rest on few studies, some of them small or testing differences in the variance rather than the mean of DNAm, and should be interpreted with caution. Pooled across studies, however, ORs for blood BrESS were close to 1 for both brain EWAS (1.03, 0.55–1.95; 19 studies) and blood EWAS of brain-related traits (0.78, 0.57–1.06; 59 studies; 1.05, 0.80–1.38, for the 43 studies of psychiatric and neurological disorders), with similar estimates for buccal BrESS (brain EWAS, 0.97, 0.53–1.79; saliva and buccal EWAS of brain-related traits, 0.93, 0.63–1.38; 7 studies) (Fig. 7A and Supplementary Fig. 6; Supplementary Table 17).

**Figure 7.**
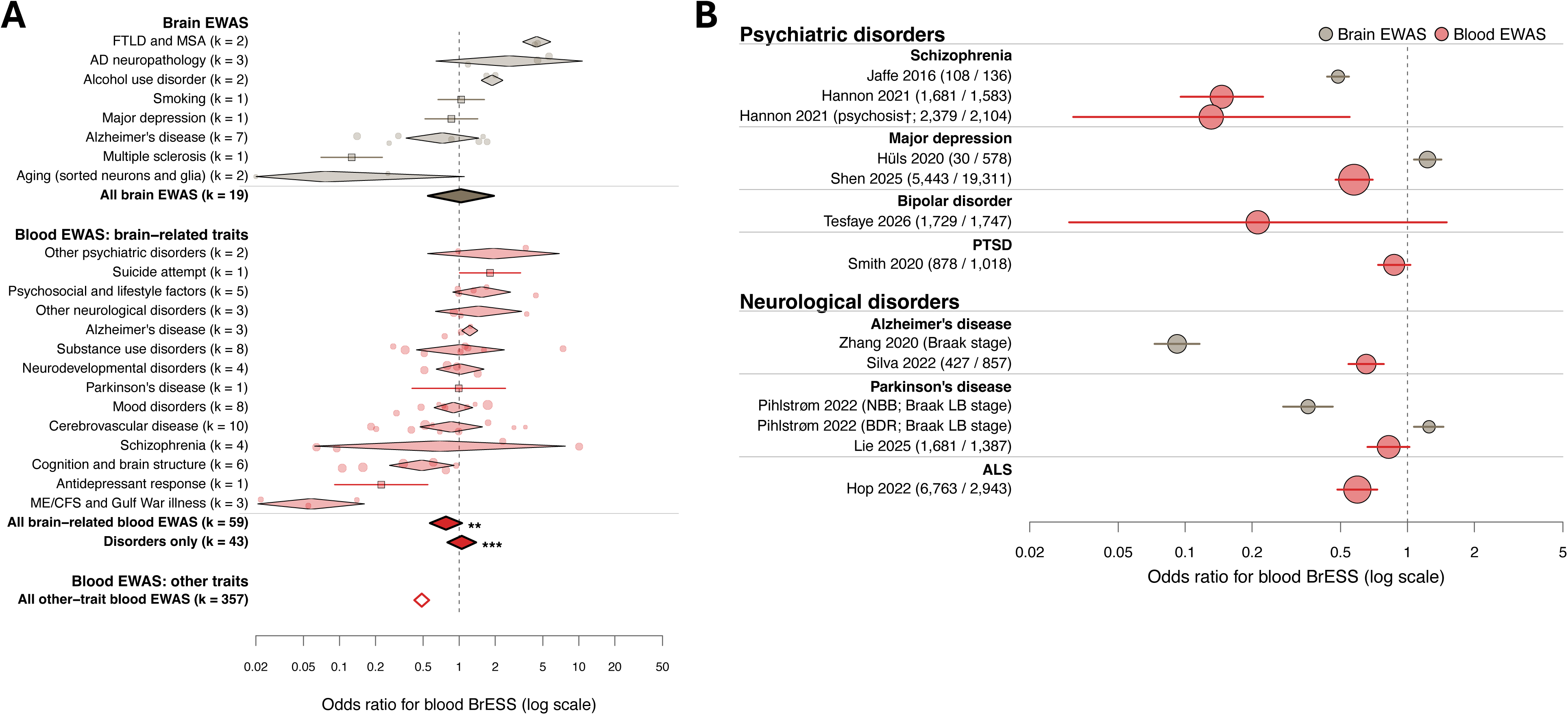
Overlap of BrESS with CpGs reported in published and large-scale EWAS. **(A)** Overlap between CpGs reported by EWAS in the EWAS Atlas and the EWAS Catalog and blood BrESS, grouped by trait within each tissue. Sections show brain EWAS, blood EWAS of brain-related traits and blood EWAS of other traits. Odds ratios are adjusted for CpG location (relation to CpG island × gene region). Small circles indicate individual studies, and their size reflects sample size on a logarithmic scale. Diamonds indicate pooled estimates for each trait group with 95% confidence intervals. Squares with horizontal lines indicate trait groups represented by a single study. Bold diamonds indicate pooled estimates across all studies in each section. The diamond labelled ‘Disorders only’ indicates the pooled estimate for studies of psychiatric and neurological disorders among the brain-related traits. The open diamond indicates the pooled estimate for blood EWAS of other traits. Asterisks indicate pooled estimates that differ from the open diamond by random-effects meta-regression (**P < 0.01; ***P < 0.001). k denotes the number of studies. AD neuropathology includes amyloid plaques, neurofibrillary tangles and Braak stage. **(B)** Overlap between disease-associated CpGs from large-scale EWAS and blood BrESS. Each row shows one EWAS. Circles indicate odds ratios, and horizontal lines indicate 95% confidence intervals. Circle size reflects study sample size on a logarithmic scale, and numbers in parentheses give cases / controls. For studies of neuropathological stage, the stage measure is shown in parentheses in place of case and control numbers. †Psychosis cases comprise patients with schizophrenia or first-episode psychosis. NBB, Netherlands Brain Bank; BDR, Brains for Dementia Research. In both panels, colors indicate the tissue of the EWAS (gray, brain; red, blood), the vertical dashed line marks an odds ratio of 1 and the x-axis is on a log scale. Pooled estimates, group comparisons and the list of studies are in Supplementary Tables 17, 18 and 19, and large-scale EWAS details and full results are in Supplementary Tables 20 and 21.

In contrast, CpGs reported by blood EWAS of traits unrelated to the brain were depleted of blood BrESS (0.49, 0.42–0.56; 357 studies). The pooled OR was therefore higher for blood EWAS of brain-related traits than for those of other traits (ratio 1.63, 95% CI 1.17–2.27, P = 0.0042), particularly for psychiatric and neurological disorders (2.10, 1.45–3.04, P = 8.4 × 10⁻⁵) (Fig. 7A; Supplementary Table 18). This contrast is consistent with the design of BrESS as sites at which peripheral DNAm reflects the brain, although it arises from the depletion of BrESS among CpGs reported for other traits rather than an enrichment among those for brain-related traits. Although saliva and buccal EWAS showed the same direction, the difference was not significant with 7 and 6 studies (ratio 2.71, P = 0.24; Supplementary Fig. 6).

### BrESS in large-scale EWAS of psychiatric and neurological disorders

Finally, we examined CpGs reported by large-scale EWAS of psychiatric and neurological disorders^4,5,9,10,38–44^ (Fig. 7B; Supplementary Tables 20 and 21). Among brain EWAS, CpGs associated with major depressive disorder and with Parkinson’s disease in one of two brain-bank cohorts (BDR) were modestly enriched in blood BrESS (OR 1.23, 95% CI 1.07–1.42; OR 1.25, 95% CI 1.07–1.45), whereas those for schizophrenia, Alzheimer’s disease and Parkinson’s disease in the other cohort (NBB) were depleted (OR 0.09–0.49). None of the blood EWAS showed enrichment (OR 0.13–0.87). When brain and blood EWAS of the same disorder were compared, overlap with blood BrESS was higher for the brain EWAS for schizophrenia (ratio of ORs 3.34, 95% CI 2.15–5.18), major depressive disorder (2.14, 1.69–2.71) and Parkinson’s disease in BDR (1.52, 1.17–1.98), but lower for Alzheimer’s disease (0.14, 0.11–0.19) and Parkinson’s disease in NBB (0.43, 0.31–0.61) (Supplementary Table 22). These ratios are relative. For schizophrenia, for example, the brain EWAS was itself depleted in blood BrESS (OR 0.49), but less so than the blood EWAS (OR 0.15).

## Discussion

In this study, we substantially increased the amount of paired brain and peripheral DNAm data from the same individuals and built three resources, which together provide an empirical basis for interpreting peripheral DNAm data in terms of their correlation with the brain. IMAGE-CpG2 showed that the pattern of brain–peripheral correlation is largely preserved across the conditions under which brain tissue is obtained. Site-level correlations were highly concordant between brain regions, including the hippocampus and amygdala, and their distributions were similar across the biopsy, autopsy and pediatric cohorts. For blood in particular, correlations differed little between the biopsy and autopsy cohorts, suggesting that estimates obtained from post-mortem brain also inform how blood relates to the living brain. These data also allowed us to revisit which peripheral tissue best serves as a surrogate for the brain. Because buccal epithelium shares an ectodermal origin with the brain, it has been suggested to be a superior surrogate to blood or saliva^13^. We found no clear difference between blood and buccal epithelium in the magnitude of their correlation with the brain (saliva could not be conclusively evaluated owing to its limited sample size). If anything, blood was more reproducible across cohorts, whereas buccal correlations differed slightly between cohorts that also differed in age composition and collection method, suggesting that buccal-specific characteristics may affect its stability as a surrogate. Expanding from IMAGE-CpG2, COMPASS provides, to our knowledge, the first meta-analytic integration of multiple cohorts for brain–peripheral DNAm correlation^14–18^, and quantifies between-cohort heterogeneity at each CpG so that single-cohort estimates can be interpreted against the consensus.

Building on these resources, the central contribution of this study is the genome-wide definition of BrESS. Although brain–peripheral correlation has previously been reported as correlation values at individual sites, no framework had defined, with an explicit and validated criterion, the full set of sites at which peripheral DNAm reliably reflects brain DNAm. BrESS are selected not by the strength of correlation in a single dataset but by out-of-sample prediction that is reproducible across cohorts differing in collection method.

Prediction at BrESS was also partly reproduced in an independent cohort. This supports the use of BrESS as a catalog, while showing that prediction at individual sites is moderate. The value of TRACE-CpG therefore lies not in replacing measurement of the brain but in the catalog of sites at which the correspondence between peripheral and brain DNAm is supported by prediction (the interpretation of pBrain values is described in Supplementary Note 2).

We next characterized BrESS against external reference data. BrESS were concentrated at CpG-island promoters that are active in both the brain and the peripheral reference epigenomes, and in genes that are expressed across tissues and serve housekeeping functions such as DNA repair, nucleic acid metabolism and the mitotic cell cycle.

Conversely, they were depleted at enhancers and at bivalent and Polycomb-repressed regions, in genes preferentially expressed in the brain or in the periphery and in genes involved in development, synaptic transmission and growth factor signaling. BrESS were enriched among CpGs with the strongest mQTL effects, but this enrichment was confined to the most extreme sites, and most BrESS were not explained by known mQTLs. This genetically driven component includes CoRSIVs, regions that vary systemically across tissues under strong genetic influence^23^, which were strongly enriched among BrESS but account for less than 1% of them. BrESS were also not enriched among CpGs at which differences in DNAm between individuals are associated with differences in the expression of nearby genes in blood. Taken together, the variation in DNAm that the periphery shares with the brain lies mainly at promoters that are active in both the brain and the periphery and belong to broadly expressed housekeeping genes, rather than at regulatory elements that differ between tissues. For most BrESS, this variation is not attributable to strong genetic effects and is not accompanied by variation in the expression of nearby genes.

Because these promoters are in the same active state in the brain and the periphery, the factors that set DNAm at these sites are likely to be shared across tissues. One possibility is that inter-individual differences at these sites arise in regulatory states established early in development, before tissues diverge, and are subsequently maintained across tissues^45^, as is thought to be the case for CoRSIVs^23^. Another is that factors acting on the whole body, such as the systemic physiological environment, affect DNAm at these promoters similarly in the brain and the periphery. In either case, BrESS capture variation in a regulatory compartment shared across tissues, rather than in the tissue-specific regulation that distinguishes the brain from other tissues.

This character of BrESS is also reflected in their relationship to the genetic risk of psychiatric and neurological disorders. SNPs near BrESS did not contribute to the heritability of any of the seven disorders. Because the baseline annotations already include promoters and transcription start sites, this test is conservative, and the absence of an effect means that BrESS carry no heritability beyond that of promoter CpGs in general, not that risk variants are absent from these promoters. The genetic risk of these disorders is thus not concentrated at BrESS, in line with the finding that most BrESS are not under strong genetic control. Beyond genetic risk, a more direct implication of this characterization concerns how BrESS relate to disease-associated DNAm. Peripheral epigenetic studies have often assumed that CpGs at which brain and peripheral DNAm correlate are where peripheral signals are most likely to be relevant to the brain. Our results qualify this assumption. Pooled across hundreds of studies in the EWAS Atlas and the EWAS Catalog, CpGs reported for brain-related traits, including psychiatric and neurological disorders, were neither enriched nor depleted for BrESS. Large-scale EWAS showed the same picture at higher resolution. Disease-associated CpGs from brain were at most modestly enriched for BrESS and were depleted for several disorders, and those from blood were consistently depleted. Accordingly, when brain and blood EWAS of the same disorder were compared, the brain EWAS overlapped more with BrESS for some disorders but not for others. This pattern is what the character of BrESS would predict. BrESS lie at promoters active across tissues, whereas disease-related variation is likely to involve tissue-specific regulation, such as that of the developmental and synaptic genes in which BrESS were depleted. Together with the genetic analyses, these results indicate that the genetic and epigenetic signals of psychiatric and neurological disorders lie largely outside the part of the epigenome that the brain shares with the periphery.

This does not make peripheral EWAS signals outside BrESS uninformative about disease. Although such signals cannot be attributed to the brain from peripheral data, they may capture disease-related processes in the peripheral tissue itself, including immune and metabolic responses, changes in cell composition and the effects of medication, smoking and other exposures. Consistent with this view, CpGs reported for traits unrelated to the brain, many of which reflect such exposures and physiological states, were depleted of BrESS. Conversely, a CpG outside BrESS should not be read as unimportant for the brain, only as lacking a reliable peripheral counterpart, because brain-specific regulation that leaves no trace in peripheral tissue lies outside a correlation-based framework by construction^46^. Correlation with the brain and association with disease are therefore distinct properties of a CpG. BrESS are best used not as a filter that selects disease signals but as a means of determining which peripheral signals can be attributed to the brain, by indicating for each CpG whether peripheral DNAm is expected to have a counterpart in the brain.

Several limitations should be considered. The brain tissue analyzed here comes from two sources, neither of which is a neutral window onto the healthy living brain. Living samples were obtained during resection for medically intractable epilepsy, whereas post-mortem samples are subject to peri-mortem influences, although the pattern of brain–peripheral correlation was similar between them. In addition, the DNAm array on which this study relies covers only a small fraction of genomic CpGs, and its probe design is biased toward gene promoters and CpG islands^47^. Although the promoter and island enrichment of BrESS likely reflects a genuine biological tendency, potential contributions from biases inherent to this measurement platform cannot be fully excluded. Characterizing the full landscape of sites at which peripheral DNAm reflects brain DNAm will require extending these resources toward whole-genome measurement, and integrating them with approaches that draw on information independent of peripheral correlation, such as prediction of DNAm from DNA sequence^48,49^. Finally, the brain and peripheral DNAm measured here are all based on bulk samples comprising multiple cell types, such as neurons and glia in the brain, leukocyte subtypes in blood and epithelial cells mixed with leukocytes in buccal samples. Because cell-type composition was not adjusted for, we cannot exclude the possibility that part of the observed correspondence reflects inter-individual variation in these proportions within each tissue^50–52^, and BrESS do not indicate in which brain cell types the shared variation arises. Addressing this will require cell-type-resolved data, such as DNAm from sorted neuronal and glial nuclei.

Despite these limitations, these resources place the study of the human brain using peripheral DNAm on firmer empirical ground. BrESS define where peripheral DNAm can be read as a reflection of brain DNAm, and show that these sites capture variation in a regulatory compartment shared across tissues rather than the tissue-specific regulation in which disease signals mostly lie. As a reference, BrESS allow peripheral DNAm studies to distinguish signals that can be attributed to the brain from those that cannot.

## Methods

### Study cohorts and participants

We assembled three newly compiled cohorts of matched brain and peripheral DNAm samples, collectively referred to as IMAGE-CpG2 (n = 148 donors): an adult surgical biopsy cohort (DB6; n = 40), an adult autopsy cohort (DB7; n = 82) and a pediatric cohort (DB8; n = 26) (Table 1). Of these, 15 donors in DB6 and 6 in DB8 (21 in total) were previously reported as part of the original IMAGE-CpG resource^16^ (GEO accession GSE111165) and re-analyzed here. The remaining donors were newly generated for this study. Up to three peripheral tissues (whole blood, buccal epithelium and saliva) were collected per donor, and the available tissues and brain regions, together with donor age and sex, are summarized in Table 1. Donors were predominantly of European ancestry, with smaller numbers of East Asian and other ancestries. The samples were collected under protocols approved by the University of Iowa Human Subjects Research Institutional Review Board and the Stanford University Institutional Review Board (#62033), with written informed consent obtained from all participants or their parents or legal representatives, as applicable.

### Sample collection

Brain tissue was obtained either at neurosurgery (biopsy donors) or at autopsy (autopsy donors), and the biopsy and autopsy portions of each cohort are indicated in Table 1.

Biopsy donors underwent neurosurgical resection for medically intractable epilepsy, and brain tissue was obtained from the resected epileptogenic lesion, so the available brain region was determined by the surgical site rather than predefined anatomically. For these donors, brain tissue was handled as previously described for the IMAGE-CpG resource^16^.

Autopsy donors were sampled at the dorsolateral prefrontal cortex, amygdala and hippocampus where available, although not all regions could be obtained from every donor. The post-mortem interval was available for 52 DB7 donors (mean 15.4 hours) and for 3 of 4 DB8 autopsy donors (mean 15.3 hours). Whole blood was collected in EDTA tubes, saliva using the Oragene kit and buccal epithelium using swabs. All samples were stored at −80 °C until DNA extraction.

### DNA extraction

Genomic DNA was extracted from brain tissue, whole blood, buccal epithelium and saliva using the MasterPure DNA extraction kit (Epicentre, MCD85201), following tissue-specific protocols. DNA quality was assessed by NanoDrop spectrophotometry, and DNA was quantified using the Qubit dsDNA Broad Range Assay Kit (Thermo Fisher Scientific, Q32850).

### DNAm profiling

For each sample, 500 ng of genomic DNA was bisulfite converted using the EZ DNA Methylation Kit (Zymo Research, D5002). Genome-wide DNAm was profiled using Illumina Infinium MethylationEPIC BeadChip arrays. Samples were assayed across EPIC v1 and v2 array versions, with the version composition differing by cohort (DB6, EPIC v1 n = 40; DB7, EPIC v1/v2/v2C n = 26/19/37; DB8, EPIC v1/v2C n = 23/3; Table 1). Here, v2C denotes a customized version of the EPIC v2 array, for which the platform annotation has been deposited in GEO (GPL34510). Samples were randomized across chips to minimize batch effects.

### Data preprocessing and quality control

Raw IDAT files were processed using a pipeline adapted from the Psychiatric Genomics Consortium (PGC) PTSD EWAS group (https://github.com/skatrinli/PGC-PTSD-EWAS-EPIC_QC), with a parallel implementation for the EPIC v2 platform (https://github.com/skatrinli/EPICv2_pipeline). Because the cohorts were assayed on multiple array versions, the EPIC v2 workflow was harmonized with that for v1 so that v1, v2 and v2C data were processed equivalently. For all versions, background and dye-bias correction was performed with the Noob method (preprocessNoob, single-sample dye correction^53^) in minfi^54^ using the appropriate EPIC manifest. Probes were removed if they had a detection P value > 0.01 or were missing in more than 10% of samples. Cross-reactive and off-target probes were removed using previously published annotations^55,56^, and for EPIC v2 and v2C, replicate probes targeting the same CpG were collapsed according to Peters et al.^56^. DNAm levels were expressed as beta values for descriptive analyses and converted to M-values (M = log2(beta/(1 − beta))) for statistical modeling.

Predicted sex was compared with recorded sex as a quality check. Discordances were typically confined to a single tissue from a donor whose other tissues matched, and sample identity across tissues from the same donor was confirmed using the genotyping (SNP) probes on the array, indicating array-level technical artifacts rather than sample mix-ups. Such samples were therefore retained. We also did not apply the control-probe-based QC-failure flagging or SNP-based outlier exclusion included in the standard pipeline, as these would have removed a substantial number of these valuable specimens. Because samples were assayed on different EPIC versions, analyses were restricted to CpG probes shared across the relevant array versions, yielding a common set of 687,873 CpG sites used in all cross-tissue analyses.

### Brain–peripheral correlation atlas (IMAGE-CpG2)

Global DNAm structure was visualized by multidimensional scaling (MDS) of genome-wide M-values, based on Euclidean distances between samples, within each cohort and in the integrated dataset. For each CpG site, Spearman’s correlation coefficient (ρ) was calculated between brain and peripheral DNAm across paired donors, following our previous study^16^. Spearman’s correlation is robust to residual technical variation, which was relevant here because no batch correction was applied to the DNAm data.

Correlations were computed both for individual brain regions and for pan-brain DNAm, defined as the within-subject average across all available brain regions in each donor. In DB7, where multiple regions were available within donors, region-specific correlations between each brain region and each peripheral tissue, as well as correlations between paired brain regions, were computed. To assess the reproducibility of site-level correlations across cohorts, CpG-wise ρ values were compared pairwise between the three newly collected cohorts for each peripheral tissue, and their agreement was quantified by Pearson correlation. To compare biopsy and autopsy collections directly, the per-CpG difference in ρ between DB6 and DB7 (Δρ = ρ_DB6 − ρ_DB7) was computed across all CpG sites for each tissue and examined both as a distribution and against the mean ρ of the two cohorts.

### Cross-cohort meta-analysis (COMPASS)

To integrate brain–peripheral correlations across cohorts, IMAGE-CpG2 (DB6, DB7 and DB8) was combined with five previously published paired brain–peripheral datasets (DB1–DB5): Hannon et al.^14^ (DB1), Edgar et al.^15^ (BECon; DB2), Braun et al.^16^ (IMAGE-CpG; DB3), Sommerer et al.^17^ (DB4) and Nishitani et al.^18^ (AMAZE-CpG2, expanded in this study; DB5) (Table 1). For each published cohort, CpG-wise Spearman correlations (ρ) between pan-brain and peripheral DNAm were obtained or recomputed on the same basis as for IMAGE-CpG2, and meta-analyses were performed separately for each peripheral tissue. Owing to differences in tissue coverage across cohorts, the contributing cohorts differed by tissue: blood (DB1–DB3 and DB5–DB8), buccal epithelium (DB4–DB8) and saliva (DB3, DB5, DB6 and DB8). For each CpG site, per-cohort Spearman ρ values were transformed using Fisher’s z transformation and combined by random-effects meta-analysis with restricted maximum likelihood (REML) estimation in metafor^57^, then back-transformed to yield a pooled correlation coefficient together with a measure of between-cohort heterogeneity (I²). The meta-analysis was restricted to CpG sites available across the contributing cohorts.

### Identification of brain epigenome shared sites and prediction of brain DNAm (TRACE-CpG)

To model the relationship between brain and peripheral DNAm, we fit a linear mixed model (LMM)^58^ at each CpG site for blood and for buccal epithelium. Brain DNAm (M-value) was modeled as a function of peripheral DNAm, age and sex as fixed effects, with donor and brain region included as random effects to account for the repeated sampling of multiple brain regions per donor:

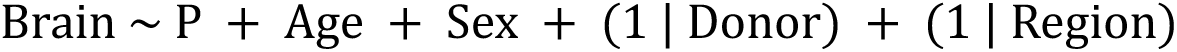

where Brain and P denote the M-values of brain and peripheral DNAm, respectively. A reduced model excluding the peripheral term was fit in the same way. Models were fit by maximum likelihood, which permits comparison of the full and reduced models that differ in their fixed-effect structure.

BrESS were identified by leave-one-dataset-out cross-validation (LODO-CV) across the three IMAGE-CpG2 cohorts (DB6, DB7 and DB8). In each of the three cycles, the full and reduced models were trained on two cohorts, and their fixed-effect coefficients were used to predict brain DNAm in the held-out cohort. Out-of-sample R² was computed as the squared correlation between predicted and measured brain DNAm across all brain samples in the held-out cohort, with each donor’s predicted value compared with each of that donor’s brain regions, and R²gain was defined as the out-of-sample R² of the full model minus that of the reduced model, that is, the gain in prediction from adding peripheral DNAm to age and sex. CpG sites with R²gain > 0.2 in at least two of the three cycles were defined as BrESS, separately for blood and buccal epithelium. Saliva was not included because DB7 had no saliva samples and DB8 had only six, which did not permit LODO-CV. The models used for prediction were then fit on all IMAGE-CpG2 data (DB6–DB8) (Supplementary Data 2). For each CpG, we computed the marginal R² (variance explained by the fixed effects) and the conditional R² (variance explained by both fixed and random effects), following the approach of Nakagawa and Schielzeth^59,60^. The variance in brain DNAm explained by peripheral DNAm (ΔR²m) was obtained as the difference in marginal R² between the full and reduced models. Using these models, brain DNAm was predicted from peripheral DNAm (pBrain) on the M-value scale as

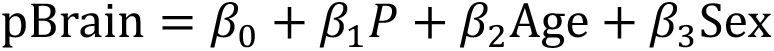

where β₀–β₃ are the coefficients estimated for the corresponding peripheral tissue, with predictions converted back to beta values where needed for visualization. Because pBrain is a linear prediction, its inter-individual variance is smaller than that of measured brain DNAm. Where pBrain is shown on the scale of brain DNAm, this variance was restored with a per-CpG scaling factor derived from IMAGE-CpG2, as described in Supplementary Note 2.

### External validation (AMAZE-CpG2)

The TRACE-CpG models fit on all IMAGE-CpG2 data were evaluated in the independent AMAZE-CpG2 cohort (DB5; n = 35; EPIC v1), which was not used for model fitting or for deriving the scaling factors. In AMAZE-CpG2, brain tissue was obtained from a single resection site per donor, which differed between donors, and was treated as pan-brain DNAm. For blood and buccal epithelium, brain DNAm was predicted from peripheral DNAm, age and sex using the corresponding TRACE-CpG model. Prediction accuracy was assessed at each BrESS by fitting an LMM of measured brain DNAm on pBrain, with resection region included as a random effect to account for differences in sampling site between donors, and r² was defined as the marginal R² of this model. The distribution of r² across BrESS was summarized for each tissue. To visualize the relationship between predicted and measured profiles, MDS was performed on the M-values at BrESS of measured peripheral DNAm, measured brain DNAm and variance-corrected pBrain, based on Euclidean distances, after adjustment for measurement batch.

### Molecular characterization of BrESS Genomic context

BrESS were compared with non-BrESS sites among autosomal CpG sites evaluated in LODO-CV for the corresponding tissue, and this background was used in all subsequent analyses unless otherwise stated. Each site was annotated with its relation to CpG island (island, shore, shelf or open sea) and its UCSC gene region from IlluminaHumanMethylationEPICanno.ilm10b4.hg19. For gene region, the first annotation was used, and sites lacking a RefSeq gene annotation were classified as intergenic. The proportions of BrESS and non-BrESS sites in each category were compared by two-sided Fisher’s exact tests. Because genomic location can confound the overlap between sets of CpG sites, overlaps between BrESS and other sets of CpG sites in the subsequent analyses were quantified as Mantel–Haenszel ORs^61^ stratified by the combination of relation to CpG island and gene region, with 95% CIs from the Robins–Breslow–Greenland variance (location-adjusted ORs)^62^. Unadjusted ORs from two-sided Fisher’s exact tests are given in the Supplementary Tables.

### Chromatin states

Each CpG site was assigned the chromatin state at its hg19 position in the Roadmap Epigenomics 15-state core model^19^ for the brain (E073, dorsolateral prefrontal cortex) and for a peripheral reference (E062, peripheral blood mononuclear cells, for blood BrESS, and E057, foreskin keratinocytes, for buccal BrESS, because no oral epithelium reference is available). The 15 states were collapsed into seven groups according to their state descriptions^19^: active TSS (TssA and TssAFlnk), transcription (TxFlnk, Tx and TxWk), enhancer (EnhG and Enh), bivalent (TssBiv, BivFlnk and EnhBiv), Polycomb (ReprPC and ReprPCWk), heterochromatin and ZNF (ZNF/Rpts and Het) and quiescent (Quies). For each of the 49 combinations of brain and peripheral state, BrESS were compared with non-BrESS sites by the location-adjusted OR.

### Gene expression

For each gene with at least five CpG sites in the EPIC UCSC_RefGene_Name annotation, the proportion of BrESS among its CpG sites was calculated, with CpG sites annotated to several genes counted for each gene. Genes were classified by the median TPM of 54 tissues in GTEx v8^20^ as housekeeping (TPM of at least 1 in all tissues), brain-enriched (mean TPM of the 13 brain regions of at least 1 and at least four times that of every other tissue), blood-enriched (TPM in whole blood of at least 1 and at least four times that of every other tissue) or epithelium-enriched (mean TPM of esophagus mucosa, minor salivary gland and skin of at least 1 and at least four times that of every other tissue). Epithelium-enriched genes were used as the periphery-enriched class for buccal BrESS. The difference in BrESS proportion between genes in each class and all other genes was estimated by linear regression with HC3 robust standard errors^63^, without adjustment and with adjustment for the genomic location of the gene’s CpG sites (in the strata used for location-adjusted ORs), log CpG number and mean expression across tissues.

### Gene Ontology analysis

The gene-level BrESS proportions were also used for Gene Ontology (GO) analysis, which proceeded from genes to GO terms to clusters of terms. First, the proportion of each gene was regressed on the genomic location of its CpG sites, as above, and on log CpG number, and genes were ranked by the residual, that is, by how much more or less BrESS they carry than expected from the location of their CpG sites. Second, preranked gene set enrichment analysis^64^ of GO biological-process terms^65^ was performed on this ranking with fgsea^66^, giving each term a normalized enrichment score (NES) that is positive when its genes carry more BrESS than expected and negative when they carry fewer. Third, terms with FDR < 0.05 were clustered by semantic similarity with simplifyEnrichment (binary cut, cutoff 0.85)^67^, separately for enriched and depleted terms, and each cluster was given a short theme name on the basis of its member terms and keyword enrichment. Clusters were defined in the blood BrESS analysis and compared by the mean and interquartile range of the NES of their member terms in blood BrESS and, for the same terms, in buccal BrESS. Clusters defined in the buccal BrESS analysis are given in Supplementary Table 12.

### mQTL

For each CpG site, the strength of mQTL association was summarized as the maximum absolute test statistic (|t| = |β|/SE) across its mQTLs, separately for cis and trans mQTLs, in two independent blood-based mQTL databases (GoDMC^21^ and EPIGEN^22^). CpG sites were ranked by |t| within each database, with sites without an mQTL ranked last, and thresholded at successive top percentiles (50%, 25%, 10%, 5%, 2%, 1%, 0.5% and 0.1%). At each threshold, blood BrESS were compared between the top-ranked and the remaining sites by the location-adjusted OR. Because GoDMC was based on the 450K array, the GoDMC analysis was restricted to CpG sites on the 450K array. CoRSIV coordinates^23^ (hg38) were converted to hg19 with liftOver, and CpG sites within CoRSIVs were compared between BrESS and non-BrESS sites in the same way.

### eQTM

Cis eQTM CpG sites at FDR < 0.05 were taken from BIOS^24^, and CpG sites in the top 10,000 cis and trans pairs, the published subset, from the Framingham Heart Study^25^, both from whole blood. Because both datasets were based on the 450K array, the background was restricted to CpG sites on the 450K array, and blood BrESS were compared between eQTM CpG sites and the remaining sites by the location-adjusted OR.

### Stratified LD score regression

GWAS summary statistics of European ancestry were obtained for seven psychiatric and neurological disorders^28–34^ and, for comparison, for educational attainment and ten non-brain traits (Supplementary Table 15). Summary statistics were processed with munge_sumstats in LDSC^68^, restricted to HapMap3 SNPs and matched to the reference by rsID. For Alzheimer’s disease, SNPs in the APOE region (chr19:43.9–46.0 Mb, hg38) were removed. For each tissue, two binary annotations were created, marking SNPs within 500 bp of any CpG site evaluated in LODO-CV (all assessed CpGs) and of any BrESS, using CpG positions in hg19. LD scores were computed from the 1000 Genomes Phase 3^69^ European reference panel with a 1-cM window. For each trait and tissue, stratified LD score regression^26^ was run with the baselineLD v2.2 model^27^, which includes promoter and transcription start site annotations, together with the two annotations, using HapMap3 SNPs excluding the MHC region as regression weights and SNPs with minor allele frequency > 5% to define the number of SNPs. The BrESS coefficient therefore reflects the contribution of BrESS beyond being an assessed CpG site and beyond the baseline annotations. Its standardized effect (τ*)^27^ was calculated as τ × number of SNPs × standard deviation of the annotation / SNP heritability. P values for τ were corrected by the Benjamini–Hochberg method across traits within each tissue. The heritability z score (SNP heritability divided by its standard error) was recorded for each trait, and partitioned estimates for traits with z < 7 were regarded as imprecise.

### Overlap with EWAS Atlas and EWAS Catalog studies

EWAS of brain, blood and saliva or buccal tissue were obtained from the EWAS Atlas^35,36^, using the CpG sites reported by each study. From the EWAS Catalog^37^, which lists associations with P < 10⁻⁴, studies of papers not included in the EWAS Atlas were added, using CpG sites that passed array-wide significance (P < 2.4 × 10⁻⁷ for the 450K array^70^ and P < 9 × 10⁻⁸ for the EPIC array^71^). Overlap with BrESS was evaluated against a background of autosomal CpG sites on both the 450K and EPIC arrays, so that studies on either array were compared on the same set of sites, and studies with at least 20 reported CpG sites within this background were included. EWAS of molecular traits (gene expression, protein, metabolite and microRNA levels) and re-analyses performed by the EWAS Catalog were not included, and two papers that also reported large-scale EWAS analyzed separately (see below) were excluded. Analyses of the same samples that differed only in covariates were combined into one estimate by averaging their log ORs and standard errors. Traits were classified by the authors into predefined categories on the basis of the trait reported by each study, and the category of each study is given in Supplementary Table 19. Brain EWAS included all traits except glioma and did not include fetal brain. Blood EWAS of brain-related traits included psychiatric disorders, neurological disorders and other brain-related traits, such as cognition and brain structure, and excluded monogenic and chromosomal neurodevelopmental syndromes. Saliva and buccal EWAS of brain-related traits included fetal alcohol spectrum disorder, early-life adversity and stress, and childhood socioeconomic status. EWAS of traits unrelated to the brain, such as smoking, aging, body mass index, asthma and type 2 diabetes, served as a comparison group for each peripheral tissue.

For each study, the location-adjusted OR for BrESS among the reported CpG sites was calculated within this background. Brain EWAS and blood EWAS were tested against blood BrESS, and brain EWAS and saliva and buccal EWAS against buccal BrESS. Study-level log ORs were pooled within each trait group by random-effects meta-analysis with restricted maximum likelihood estimation in metafor^57^ (rma.mv), with random effects for paper and for study within paper. Pooled ORs were compared between groups by random-effects meta-regression with group as a moderator, using the same random effects. Unadjusted ORs were calculated from the 2 × 2 table summed over strata, with 95% CIs based on the standard error of the log OR, and pooled in the same way.

### Large-scale EWAS of psychiatric and neurological disorders

Large-scale EWAS of psychiatric and neurological disorders in brain and blood with publicly available CpG-level results^4,5,9,10,38–44^ were analyzed separately from the EWAS Atlas and EWAS Catalog studies (Supplementary Table 20). When only a list of significant CpG sites was released, or when more than 1,000 significant CpG sites were reported, the reported set was used. Otherwise, the 1,000 CpG sites with the smallest P values in the published summary statistics were used. For Hüls et al.^5^, only CpG sites with P < 0.01 were released, and the 1,000 with the smallest P values among them were used. The background was autosomal CpG sites on both the 450K and EPIC arrays, further restricted, for EWAS with full summary statistics, to sites tested in that EWAS. Overlap with blood BrESS was quantified by the location-adjusted OR, with unadjusted ORs from two-sided Fisher’s exact tests (results for buccal BrESS are in Supplementary Table 21). To compare how disease-associated CpG sites from brain and from blood overlap with blood BrESS, the OR of the brain EWAS was divided by that of the blood EWAS for disorders with both.

For each ratio, the standard errors of the two log ORs (brain and blood) were obtained from their 95% CIs, and the CI and P value of the ratio were calculated from the difference between the two log ORs, treating them as independent. These CIs reflect the number of CpG sites in each EWAS set and do not include the sampling variation of the EWAS themselves. For Parkinson’s disease, the two brain EWAS, from two brain banks, were each compared with the same blood EWAS.

### Web resources

All four resources are available as interactive web tools through a single portal (https://epigen.shinozakilab.net): the IMAGE-CpG2 correlation atlas (https://epigen.shinozakilab.net/image2.html), the COMPASS meta-analysis platform (https://epigen.shinozakilab.net/compass.html), the TRACE-CpG tool for predicting brain DNAm at BrESS (https://epigen.shinozakilab.net/trace.html) and the AMAZE-CpG2 atlas (https://epigen.shinozakilab.net/amaze2.html).

## Data availability

The DNAm data generated in this study (DB6–DB8) have been deposited in GEO as a SuperSeries (GSE338412), comprising SubSeries GSE338405, GSE338406, GSE338407, GSE338408, GSE338409 and GSE338410. The custom EPIC v2 platform is available under accession GPL34510. The expanded AMAZE-CpG2 dataset (DB5) has been deposited under accession GSE338391. The original AMAZE-CpG is available under GSE214901 and the original IMAGE-CpG under GSE111165. Publicly available DNAm datasets analyzed in this study include GSE59685, GSE95049, GSE74193 and GSE152026. Genome-wide buccal–brain DNAm correlation results used as DB4 are available at http://www.liga.uni-luebeck.de/buccal_brain_correlation_results/. LODO-CV results for all CpG sites and the TRACE-CpG models for BrESS are provided in Supplementary Data 1 and 2. GWAS and large-scale EWAS summary statistics were obtained from the sources listed in Supplementary Tables 15 and 20. EWAS studies were obtained from the EWAS Atlas and the EWAS Catalog, chromatin states from the Roadmap Epigenomics Project, gene expression from GTEx v8, mQTLs from GoDMC and EPIGEN, and eQTMs from BIOS and the Framingham Heart Study. The LD score reference data (1000 Genomes Phase 3, HapMap3 SNPs and baselineLD v2.2) were those distributed with LDSC.

## Code availability

Custom code for TRACE-CpG, including scripts for the identification of BrESS by LODO-CV and for the prediction of brain DNAm (pBrain) at BrESS, is available at https://github.com/snishit/TRACE-CpG.

## Acknowledgements

We thank collaborators in the departments of neurosurgery and pathology at the University of Iowa, Stanford University, the University of Fukui and Yamaguchi University for their assistance with neurosurgical procedures and autopsy brain tissue sampling.

## Author Contributions

Conceptualization: S.N., M.P.S., G.S. Methodology: S.N., S.H., A.S., A.K., T.W., G.S. Software: S.N., A.S., A.K., G.S. Formal analysis: S.N. Investigation: S.N., T.I., N.J.P., M.C., M.I., Y.H., T.Yu., S.Ka., H.A., T.K., H.I., S.No., K.Y., B.A., I.G., R.S., H.D.N., N.G., E.G., A.Y., S.Ku., T.X.F., P.R.B., T.N., T.Yn., K.K., G.A.G., J.E.H., M.A.H., H.K. Resources: S.N., M.C., M.I., Y.H., T.Yu., S.Ka., H.A., T.K., H.I., S.No., K.K., G.A.G., J.E.H., M.A.H., H.K., G.S. Data curation: S.N., A.Y., P.R.B., G.S. Validation: S.H., T.W. Visualization: S.N., A.S., A.K. Supervision: T.O., J.B.P., M.P.S., G.S. Project administration: S.N., G.S. Funding acquisition: S.N., M.I., T.X.F., A.T., G.S. Writing — original draft: S.N., M.P.S., G.S. Writing — review & editing: S.N., S.H., T.I., A.S., N.J.P., A.K., M.C., T.W., M.I., Y.H., T.Yu., S.Ka., H.A., T.K., H.I., S.No., K.Y., B.A., I.G., R.S., H.D.N., N.G., E.G., A.Y., S.Ku., T.X.F., P.R.B., T.N., T.Yn., A.T., K.K., T.O., G.A.G., J.E.H., M.A.H., H.K., J.B.P., M.P.S., G.S.

## Additional information

Correspondence and requests for materials should be addressed to G.S., S.N., or M.P.S.

