## Supplementary Information (Supplementary Tables 1-7, Supplementary Figures 1-6, Supplementary Notes 1-2 and Supplementary Data legends) for "A genome-wide catalog of sites where peripheral DNA methylation reflects the brain"

**Supplementary Table 1. Within-subject DNAm correlations between brain regions in DB7 (adult autopsy).**

| Pair | *n* | Mean ρ | Median ρ | SD | IQR | % ρ>0.5 | % ρ>0.8 | % negative |
|---|---|---|---|---|---|---|---|---|
| PFC vs HIPP | 43 | 0.376 | 0.368 | 0.239 | [0.204, 0.549] | 30.9 | 4.1 | 5.4 |
| PFC vs AMY | 43 | 0.372 | 0.375 | 0.258 | [0.185, 0.568] | 33.2 | 4.3 | 8.4 |
| HIPP vs AMY | 43 | 0.380 | 0.373 | 0.235 | [0.205, 0.554] | 31.9 | 3.6 | 4.9 |
| PFC vs HIPP (sensitivity) | 78 | 0.451 | 0.437 | 0.228 | [0.273, 0.646] | 41.9 | 5.4 | 1.4 |

At each of 687,873 CpG sites, Spearman correlation (ρ) was computed between DNAm in two brain regions across individuals in DB7 (adult autopsy) with both regions available, and the table summarizes the distribution of ρ across sites. The sensitivity analysis used all individuals with both PFC and HIPP available, without requiring AMY. n, number of paired individuals; IQR, interquartile range of ρ; PFC, prefrontal cortex; HIPP, hippocampus; AMY, amygdala.

**Supplementary Table 2. Pan-brain versus peripheral tissue DNAm correlation across DB6, DB7, DB8 and the integrated IMAGE-CpG2 dataset.**

| DB | Tissue | *n* | Mean ρ | Median ρ | SD | IQR | ρ>0.2 | ρ>0.5 | ρ>0.8 | ρ<0 |
|---|---|---|---|---|---|---|---|---|---|---|
| DB6 | Blood | 40 | 0.304 | 0.313 | 0.264 | [0.108, 0.512] | 64.2 | 26.5 | 1.4 | 14.2 |
| (Adult biopsy) | Buccal | 32 | 0.232 | 0.229 | 0.272 | [0.038, 0.429] | 54.1 | 18.0 | 1.1 | 20.9 |
| | Saliva | 32 | 0.239 | 0.230 | 0.229 | [0.072, 0.398] | 54.9 | 13.7 | 0.7 | 15.4 |
| DB7 | Blood | 72 | 0.328 | 0.325 | 0.257 | [0.136, 0.522] | 66.8 | 27.5 | 2.3 | 10.5 |
| (Adult autopsy) | Buccal | 81 | 0.336 | 0.312 | 0.268 | [0.118, 0.563] | 63.5 | 31.6 | 3.0 | 10.2 |
| DB8 | Blood | 23 | 0.301 | 0.314 | 0.253 | [0.123, 0.493] | 65.5 | 24.0 | 1.1 | 12.9 |
| (Pediatric) | Buccal | 10 | 0.244 | 0.273 | 0.372 | [-0.018, 0.539] | 56.3 | 29.4 | 4.5 | 26.7 |
| | Saliva | 6* | 0.053 | 0.086 | 0.432 | [-0.257, 0.371] | 36.8 | 16.6 | 3.1 | 45.0 |
| IMAGE-CpG2 | Blood | 135 | 0.337 | 0.346 | 0.238 | [0.159, 0.520] | 69.9 | 28.1 | 1.4 | 8.8 |
| (Integrated) | Buccal | 123 | 0.334 | 0.329 | 0.256 | [0.119, 0.547] | 64.8 | 30.7 | 2.1 | 9.9 |
| | Saliva | 38 | 0.223 | 0.215 | 0.212 | [0.071, 0.367] | 52.7 | 10.1 | 0.4 | 14.7 |

At each of 687,873 CpG sites, Spearman correlation (ρ) was computed between pan-brain DNAm (averaged across all available brain regions within each subject) and peripheral tissue DNAm across paired subjects, within each DB separately and in the integrated IMAGE-CpG2 dataset, and the table summarizes the distribution of ρ across sites. The $\rho > 0.2$, $\rho > 0.5$, $\rho > 0.8$ and $\rho < 0$ columns show the percentage of CpG sites meeting each threshold. DB7 did not include saliva samples. n, number of paired subjects; IQR, interquartile range of ρ. *DB8 × saliva results ($n = 6$) should be interpreted with caution because of the small sample size.

**Supplementary Table 3. Identification of brain epigenome shared sites (BrESS) by leave-one-dataset-out cross-validation (LODO-CV).**

| | $R^2$gain, median [IQR] | # with $R^2$gain > 0.2 (%) |
|---|---|---|
| Blood model | | |
| Cycle 1 (DB6 + DB7 → DB8) | 0.041 [−0.005, 0.202] | 173,124 (25.2%) |
| Cycle 2 (DB6 + DB8 → DB7) | 0.030 [0.000, 0.132] | 120,356 (17.5%) |
| Cycle 3 (DB7 + DB8 → DB6) | 0.026 [0.000, 0.127] | 104,378 (15.2%) |
| Passed in 3/3 cycles | – | 43,277 (6.3%) |
| Passed in 2/3 cycles | – | 77,890 (11.3%) |
| BrESS (passed in ≥ 2/3 cycles) | – | 121,167 (17.6%) |
| Buccal epithelium model | | |
| Cycle 1 (DB6 + DB7 → DB8) | 0.006 [−0.060, 0.134] | 132,560 (19.3%) |
| Cycle 2 (DB6 + DB8 → DB7) | 0.021 [0.000, 0.166] | 149,290 (21.7%) |
| Cycle 3 (DB7 + DB8 → DB6) | 0.010 [−0.001, 0.079] | 76,627 (11.1%) |
| Passed in 3/3 cycles | – | 22,616 (3.3%) |
| Passed in 2/3 cycles | – | 86,769 (12.6%) |
| BrESS (passed in ≥ 2/3 cycles) | – | 109,385 (15.9%) |

In each LODO-CV cycle, models were trained on two of the three IMAGE-CpG2 datasets (DB6, adult biopsy; DB7, adult autopsy; DB8, pediatric) and tested on the remaining one. For each CpG site, brain DNAm was predicted from a model with age and sex, with and without peripheral DNAm. $R^2$gain is the increase in out-of-sample $R^2$ for brain DNAm when peripheral DNAm is added. BrESS were defined as sites with $R^2$gain > 0.2 in at least two of the three cycles, separately for blood and buccal epithelium. In total, 76,208 sites were BrESS in both tissues (common BrESS). Percentages are relative to all CpG sites analyzed (687,873 sites shared between EPIC v1 and EPIC v2). Saliva was not included in LODO-CV because too few datasets were available for cross-validation. Per-site values are provided in Supplementary Data 1. IQR, interquartile range; –, not applicable (counts across cycles).

**Supplementary Table 4. Model fit in the discovery data and external validation of TRACE-CpG prediction models at BrESS.**

| | Blood | Buccal |
|---|---|---|
| Full-data model (IMAGE-CpG2) | | |
| Subjects with paired brain and peripheral samples, *n* | 135 | 123 |
| # of BrESS | 121,167 | 109,385 |
| $R^2$c, median [IQR] | 0.79 [0.71, 0.84] | 0.81 [0.75, 0.86] |
| $R^2$m, median [IQR] | 0.37 [0.27, 0.49] | 0.46 [0.36, 0.57] |
| $\Delta R^2$m, median [IQR] | 0.33 [0.24, 0.44] | 0.43 [0.33, 0.54] |
| # with $\Delta R^2$m > 0.2 (%) | 102,501 (84.6%) | 106,343 (97.2%) |
| External validation (AMAZE-CpG2, *n* = 35) | | |
| $r^2$, median [IQR] | 0.24 [0.10, 0.44] | 0.20 [0.07, 0.38] |
| # with $r^2$ > 0.2 / # tested (%) | 69,031 / 121,140 (57.0%) | 54,516 / 109,354 (49.9%) |

Full-data model: for each BrESS, brain DNAm (M-value) was modeled on all IMAGE-CpG2 data (DB6–DB8) with a linear mixed model, brain ~ P + Age + Sex + (1 | Donor) + (1 | Region), fit by maximum likelihood. External validation: brain DNAm was predicted (pBrain) from peripheral DNAm, age and sex in AMAZE-CpG2 (DB5, 35 subjects, EPIC v1), an independent dataset that was not used for model fitting. $r^2$ was computed for BrESS measured in AMAZE-CpG2 (blood, 121,140 of 121,167; buccal epithelium, 109,354 of 109,385). Model coefficients are provided in Supplementary Data 2. $R^2$c, conditional $R^2$, variance explained by fixed and random effects; $R^2$m, marginal $R^2$, variance explained by fixed effects (peripheral DNAm, age and sex); $\Delta R^2$m, $R^2$m of the full model minus $R^2$m of the reduced model with age and sex only, that is, the in-sample variance explained by peripheral DNAm; $r^2$, per-site marginal $R^2$ of a linear mixed model of measured brain DNAm on pBrain with brain region as a random effect; IQR, interquartile range.

### Supplementary Table 5. Subject and brain sample characteristics of AMAZE-CpG2 samples.

| | Clinical demographics | | | | | | Brain tissue | | MNI coordinate | | | Day(s) after surgery | |
|---|---|---|---|---|---|---|---|---|---|---|---|---|---|
| ID | Sex | Age (y) | Height (cm) | Weight (kg) | BMI | Diagnosis | Resection site | Hemisphere | X | Y | Z | Saliva | Buccal |
| 1 | male | 69 | 160 | 37.5 | 14.6 | primary central nervous system vasculitis | dlPFC | right | 38 | 34 | 44 | 7 | 7 |
| 2 | female | 59 | 150 | 45.5 | 20.2 | falx meningioma | frontal pole | right | 42 | 41 | 18 | 3 | 3 |
| 3 | male | 60 | 164.5 | 66.8 | 24.7 | internal carotid artery-posterior communicating artery aneurysm | temporal pole | left | -50 | 14 | -18 | 11 | 11 |
| 4 | female | 73 | 157 | 40 | 16.2 | metastatic brain tumor (breast cancer) | visual cortex | left | -45 | -85 | 18 | 7 | 7 |
| 6 | male | 61 | 164 | 69.3 | 25.8 | anterior communicating artery aneurysm | straight gyrus | left | -8 | 24 | -19 | 16 | 8 |
| 7 | male | 69 | 169 | 57 | 20.0 | internal carotid artery stenosis | temporal pole | right | 38 | 13 | -29 | 9 | 9 |
| 8 | male | 67 | 168 | 65 | 23.0 | frontal lobe tumor | dlPFC | right | 29 | 51 | 34 | 15 | 15 |
| 9 | female | 45 | 175.5 | 83 | 26.9 | internal carotid artery stenosis | temporal pole | left | -38 | 16 | -26 | 7 | 7 |
| 10 | female | 70 | 155 | 70 | 29.1 | anterior communicating artery aneurysm | orbitofrontal cortex | right | 9 | 16 | -22 | 6 | 6 |
| 11 | female | 73 | 152 | 44.1 | 19.1 | internal carotid artery-posterior communicating artery aneurysm | orbitofrontal cortex | left | -18 | 28 | -22 | 15 | 9 |
| 12 | male | 60 | 172 | 63.9 | 21.6 | tentorial meningioma | middle temporal gyrus | right | 63 | -32 | -10 | 27 | 27 |
| 13 | male | 50 | 170 | 61 | 21.1 | internal carotid artery-posterior communicating artery aneurysm | insula | right | 28 | 14 | -17 | 15 | 15 |
| 14 | female | 60 | 160.5 | 49.7 | 19.3 | middle cerebral artery aneurysm | dorsal entorhinal cortex | right | 34 | 4 | -17 | 12 | 12 |
| 15 | female | 13 | 160 | 50 | 19.5 | temporal lobe epilepsy, glioma | temporal pole | left | -60 | 2 | -26 | 11 | 42 |
| 16 | male | 62 | 185 | 93.4 | 27.3 | frontal lobe meningioma, epilepsy | frontal eye field | right | 14 | 32 | 47 | 13 | 13 |
| 17 | male | 28 | 171 | 60 | 20.5 | temporal lobe epilepsy | middle temporal gyrus | right | 63 | -9 | -24 | 14 | 14 |
| 18 | male | 67 | 170 | 53.3 | 18.4 | internal carotid artery stenosis | straight gyrus | right | 7 | 18 | -16 | 10 | 10 |
| 19 | female | 51 | 162 | 55 | 21.0 | subarachnoid hemorrhage, moyamoya disease | angular gyrus | right | 36 | -80 | 45 | 11 | 11 |
| 20 | female | 73 | 152.5 | 45.5 | 19.6 | internal carotid-posterior communicating aneurysm | temporal pole | right | 33 | 8 | -33 | 0 | 6 |
| 21 | male | 80 | 162 | 51.3 | 19.5 | meningioma | frontal eye field | left | -28 | 51 | 18 | 8 | 8 |
| 22 | female | 11 | 156.6 | 47.1 | 19.2 | temporal lobe epilepsy | middle temporal gyrus | right | 68 | -28 | -11 | 10 | 10 |
| 23 | female | 65 | 148.5 | 61.1 | 27.7 | meningioma | orbitofrontal cortex | right | 6 | 51 | -28 | 7 | 7 |
| 24 | male | 54 | 169 | 84.6 | 29.6 | subependymoma | supplementary motor area | left | -25 | 18 | 65 | 48 | 48 |
| 25 | female | 81 | 149 | 57.3 | 25.8 | unruptured intracranial aneurysm | superior temporal gyrus | right | 63 | -18 | -1 | 10 | 10 |
| 26 | female | 82 | 149.9 | 38.4 | 17.1 | meningioma | temporal pole | left | -44 | 17 | -27 | 8 | 8 |
| 27 | female | 46 | 160 | 66.4 | 25.9 | meningioma | frontal eye field | left | -45 | 14 | 45 | 4 | 4 |
| 28 | female | 68 | 147 | 51.1 | 23.6 | unruptured intracranial aneurysm | temporal pole | right | 50 | 16 | -16 | 6 | 6 |
| 29 | male | 73 | 171.6 | 61.4 | 20.9 | unruptured intracranial aneurysm | hippocampus | right | 18 | -8 | -16 | 9 | 9 |
| 30 | female | 39 | 179.8 | 60 | 28.6 | meningioma | temporal pole | right | 37 | 18 | -28 | 14 | 14 |
| 31 | female | 74 | 152 | 49.7 | 21.5 | unruptured intracranial aneurysm | temporal lobe | right | 54 | 4 | -8 | 8 | 8 |
| 33 | male | 64 | 165 | 70.3 | 25.8 | unruptured intracranial aneurysm | insula | left | -34 | 8 | -12 | 10 | 10 |
| 34 | female | 69 | 155.1 | 55.1 | 22.9 | unruptured intracranial aneurysm | frontal lobe | right | 46 | 24 | -4 | 6 | 6 |
| 35 | male | 55 | 174 | 77.3 | 25.5 | unruptured intracranial aneurysm | gyrus rectus | left | -4 | 20 | -20 | 7 | 7 |
| 36 | male | 53 | 170.2 | 65.5 | 22.6 | unruptured intracranial aneurysm | gyrus rectus | left | -6 | 12 | -18 | 2 | 2 |
| 37 | female | 75 | 145 | 46.7 | 22.2 | unruptured intracranial aneurysm | temporal pole | left | -48 | 4 | -12 | 6 | 6 |

Characteristics of all 35 AMAZE-CpG2 (DB5) participants. The first 19 were previously published[18] and the remaining 16 are newly reported here. MNI coordinates (X, Y, Z) are in Montreal Neurological Institute space. Day(s) after surgery, interval between brain resection and saliva or buccal collection; BMI, body mass index, weight (kg) / height (m)$^2$.

**Supplementary Table 6. Brain versus peripheral tissue DNAm correlation in AMAZE-CpG2 (DB5).**

| DB | Tissue | *n* | Mean ρ | Median ρ | SD | IQR | ρ>0.2 | ρ>0.5 | ρ>0.8 | ρ<0 |
|---|---|---|---|---|---|---|---|---|---|---|
| AMAZE-CpG2 (DB5) | Blood | 35 | 0.275 | 0.255 | 0.283 | [0.070, 0.477] | 57.5 | 23.0 | 3.2 | 17.0 |
| | Buccal | 35 | 0.211 | 0.187 | 0.255 | [0.026, 0.384] | 48.0 | 14.7 | 1.3 | 21.3 |
| | Saliva | 35 | 0.233 | 0.210 | 0.261 | [0.042, 0.415] | 51.4 | 17.4 | 1.6 | 19.4 |

At each of 820,406 CpG sites, Spearman correlation (ρ) was computed between brain and peripheral tissue DNAm across 35 paired subjects, and the table summarizes the distribution of ρ across sites. The ρ > 0.2, ρ > 0.5, ρ > 0.8 and ρ < 0 columns show the percentage of CpG sites meeting each threshold. n, number of paired subjects; IQR, interquartile range of ρ.

**Supplementary Table 7. Evaluation of variance-correction methods for TRACE-CpG predicted brain DNAm in AMAZE-CpG2 (DB5).**

| Method | Scaling factor | Tissue | Variance ratio (median) | Variance ratio (IQR) | $r$ (median) | RMSE (median) |
|---|---|---|---|---|---|---|
| None | 1 | Blood | 0.316 | 0.178–0.494 | 0.501 | 0.293 |
| | | Buccal | 0.388 | 0.243–0.559 | 0.452 | 0.291 |
| User-variance | $\sqrt{(\sigma^2_{brain} / \sigma^2_{pBrain, user})}$ | Blood | 2.089 | 1.396–3.196 | 0.501 | 0.419 |
| | | Buccal | 2.460 | 1.557–4.005 | 0.452 | 0.458 |
| Fixed-variance | $\sqrt{(\sigma^2_{brain} / \sigma^2_{pBrain, discovery})}$ | Blood | 0.729 | 0.490–1.008 | 0.501 | 0.309 |
| | | Buccal | 0.764 | 0.533–1.021 | 0.452 | 0.313 |

Corrected pBrain = mean(pBrain) + (pBrain − mean(pBrain)) × scaling factor, with the mean taken per CpG across the samples being predicted. A variance ratio of 1 indicates full restoration of brain-level variance. Metrics are medians across BrESS (blood, 121,140; buccal epithelium, 109,354 CpGs). *r* is identical across methods because the correction rescales values around their per-CpG mean and does not change cross-subject ordering. The Fixed-variance method is used in TRACE-CpG because it does not require any brain data from the user. The scaling factors are provided in Supplementary Data 2. pBrain, predicted brain DNAm; Variance ratio, variance of corrected pBrain divided by the variance of measured AMAZE-CpG2 brain DNAm (M-value, brain regions averaged within each subject); *r*, per-CpG correlation between corrected pBrain and measured brain across subjects; RMSE, root-mean-square error (M-value); $\sigma^2_{brain}$, variance of measured brain DNAm in the discovery (IMAGE-CpG2) data; $\sigma^2_{pBrain}$, variance of pBrain, computed either from the user's own data (User-variance, here AMAZE-CpG2) or fixed from the discovery predictions (Fixed-variance).

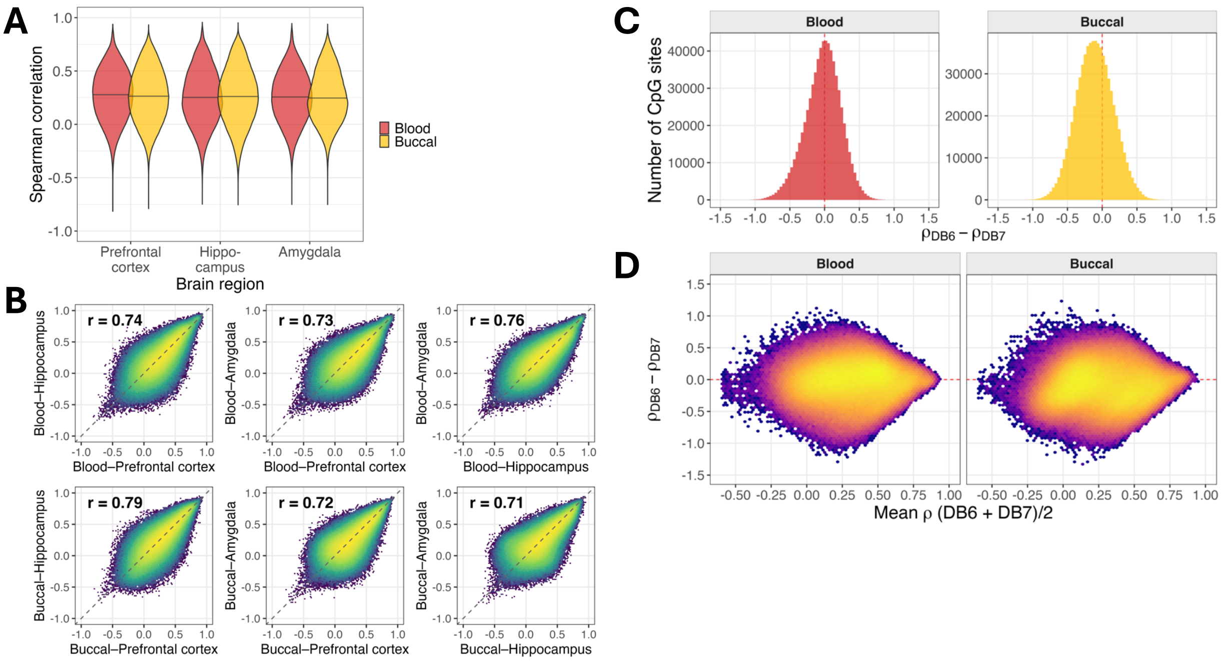


**Supplementary Figure 1. Brain–peripheral DNAm correlations: regional consistency and agreement between biopsy and autopsy cohorts.**

**(A)** Within-subject brain–peripheral DNAm correlations across brain regions in DB7 autopsy samples. Distribution of Spearman correlations between brain and peripheral tissue DNAm for each brain region (prefrontal cortex, hippocampus and amygdala) and peripheral tissue (blood and buccal epithelium), computed per CpG across 687,873 sites (n = 43 individuals with all three brain regions). Horizontal lines within violins indicate the 25th, 50th and 75th percentiles. Sensitivity analysis restricted to prefrontal cortex and hippocampus (n = 69–77) yielded consistent results (median ρ = 0.26–0.28). Inter-regional correlations are reported in Supplementary Table 1. **(B)** Concordance of per-CpG brain–peripheral correlations between brain regions. For each peripheral tissue (blood, top row; buccal epithelium, bottom row), the per-CpG Spearman correlation with one brain region is plotted against that with another, for all three region pairs (prefrontal cortex, hippocampus amygdala) across 687,873 CpG sites. Each hexagonal bin represents the density of CpG sites; the dashed line indicates the identity (y = x). The value r in each panel is the Pearson correlation between the two correlation vectors. **(C)** Histograms of the per-CpG difference in Spearman ρ (Δρ = ρ_DB6 − ρ_DB7) between DB6 (adult biopsy) and DB7 (adult autopsy) cohorts for blood (left) and buccal epithelium (right), across 687,873 CpG sites. Red dashed line indicates Δρ = 0. **(D)** Bland–Altman-style plots of Δρ (y-axis) against the mean ρ across the two cohorts ((ρ_DB6 + ρ_DB7) / 2; x-axis) for blood (left) and buccal epithelium (right). Each hexagonal bin represents the density of CpG sites. Red dashed line indicates Δρ = 0.

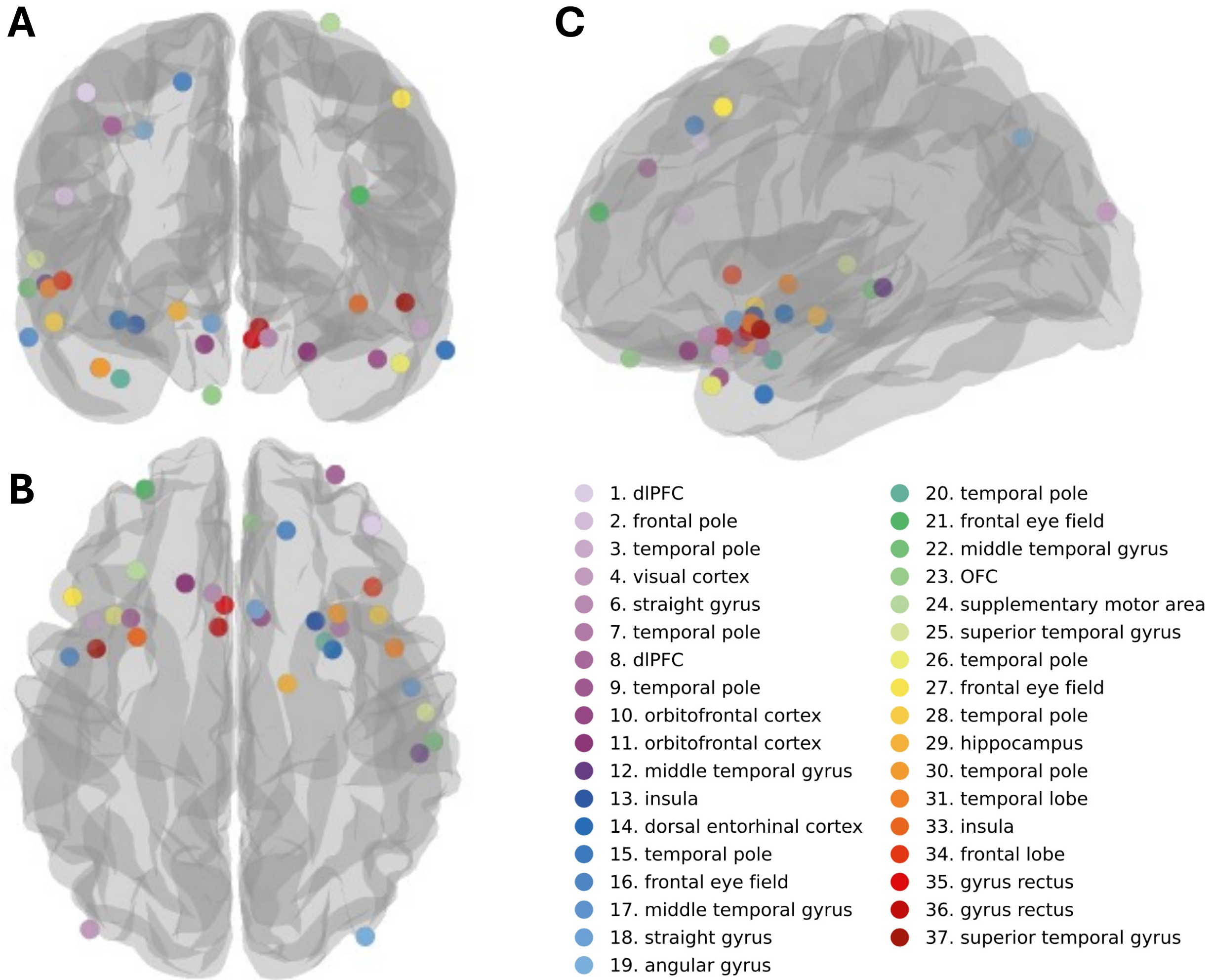


**Supplementary Figure 2. Location of each resected brain tissue according to the recorded Montreal Neurological Institute (MNI) coordinates. (A)** front, **(B)** top and **(C)** left-side views. Each sample is represented by a colored circle.

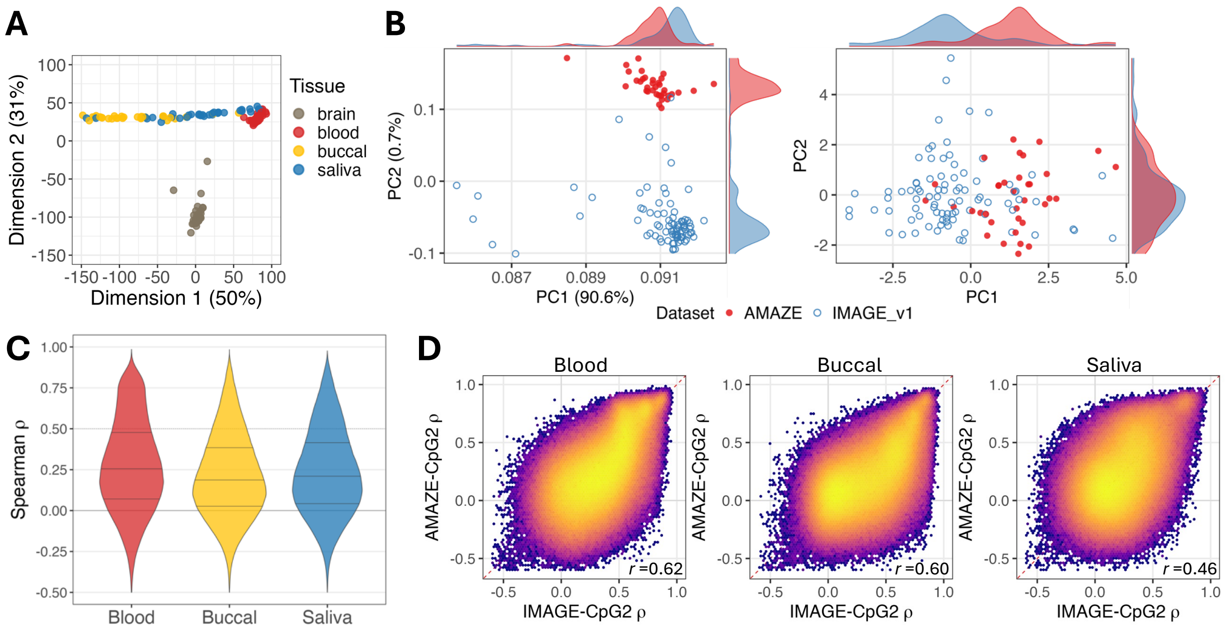


**Supplementary Figure 3. Characterization of AMAZE-CpG2 (DB5).**
**(A)** Multidimensional scaling (MDS) plot of genome-wide DNAm profiles for AMAZE-CpG2 (*n* = 35 subjects with all four tissues; 140 samples total). Each point represents a sample, colored by tissue type. Percentage of variance explained by each dimension is shown in parentheses. **(B)** Population stratification analysis restricted to EPIC v1 array samples (AMAZE-CpG2 *n* = 35; IMAGE-CpG2 v1 subset with blood DNAm, *n* = 86) to ensure probe compatibility, because EPIC v2 was redesigned with substantial removal of SNP-overlapping probes used for ancestry inference. Left: Barfield approach (the method used in the original AMAZE-CpG study; *n* = 5,825 SNP-CpGs; percent variance explained indicated). Right: EpiAnceR+ approach (*n* = 3,596 SNP-CpGs; axes reflect residualized variance after adjustment for cell composition, sex, age, and technical variation). Both methods reveal clear ancestry separation between AMAZE-CpG2 (filled red, Japanese) and IMAGE-CpG2 v1 (open blue, predominantly European). Marginal density plots shown along each axis. **(C)** Distribution of CpG-wise brain–peripheral DNAm correlations (Spearman ρ) in AMAZE-CpG2 for blood, buccal epithelium and saliva. For each tissue, ρ was computed per CpG site between brain and peripheral DNAm across the 35 paired subjects, over 820,406 CpG sites. Horizontal lines within each violin indicate the 25th, 50th and 75th percentiles. Summary statistics are in Supplementary Table 6. **(D)** Pairwise concordance of CpG-wise ρ values between the integrated IMAGE-CpG2 dataset (x-axis) and AMAZE-CpG2 (y-axis) for blood, buccal epithelium and saliva, across 687,798 CpG sites common to both datasets. Red dashed line indicates y = x. Pearson r in the bottom-right corner quantifies the agreement of ρ values between the two independent datasets.

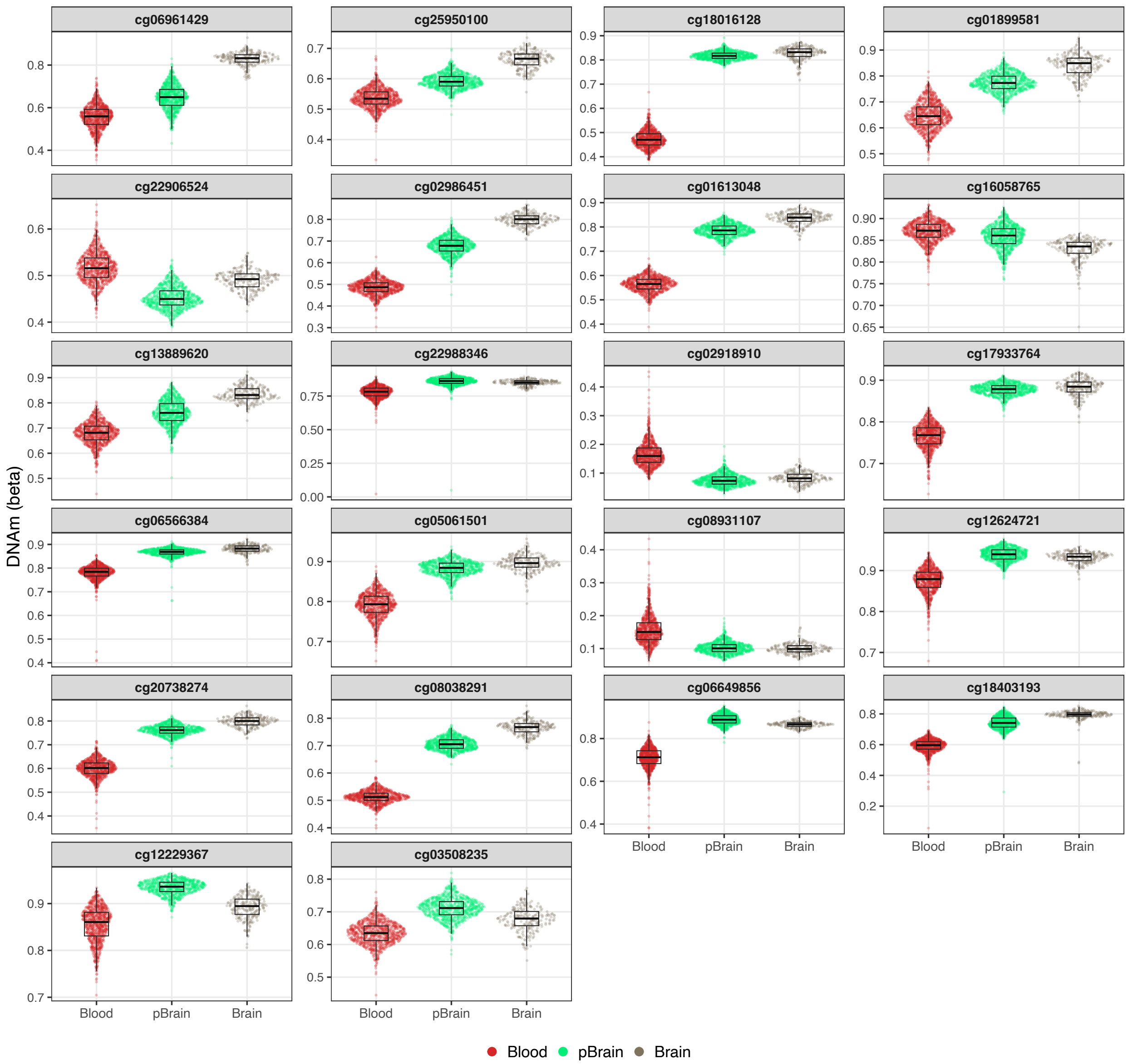


**Supplementary Figure 4. Blood, TRACE-predicted brain (pBrain) and measured brain DNAm at 22 representative CpGs.** Blood and pBrain, GSE152026 (*n* = 934); Brain, GSE74193 (*n* = 244). Each point is one sample. Boxes show the median and IQR, and whiskers extend to 1.5 × IQR (details in Supplementary Note 2).

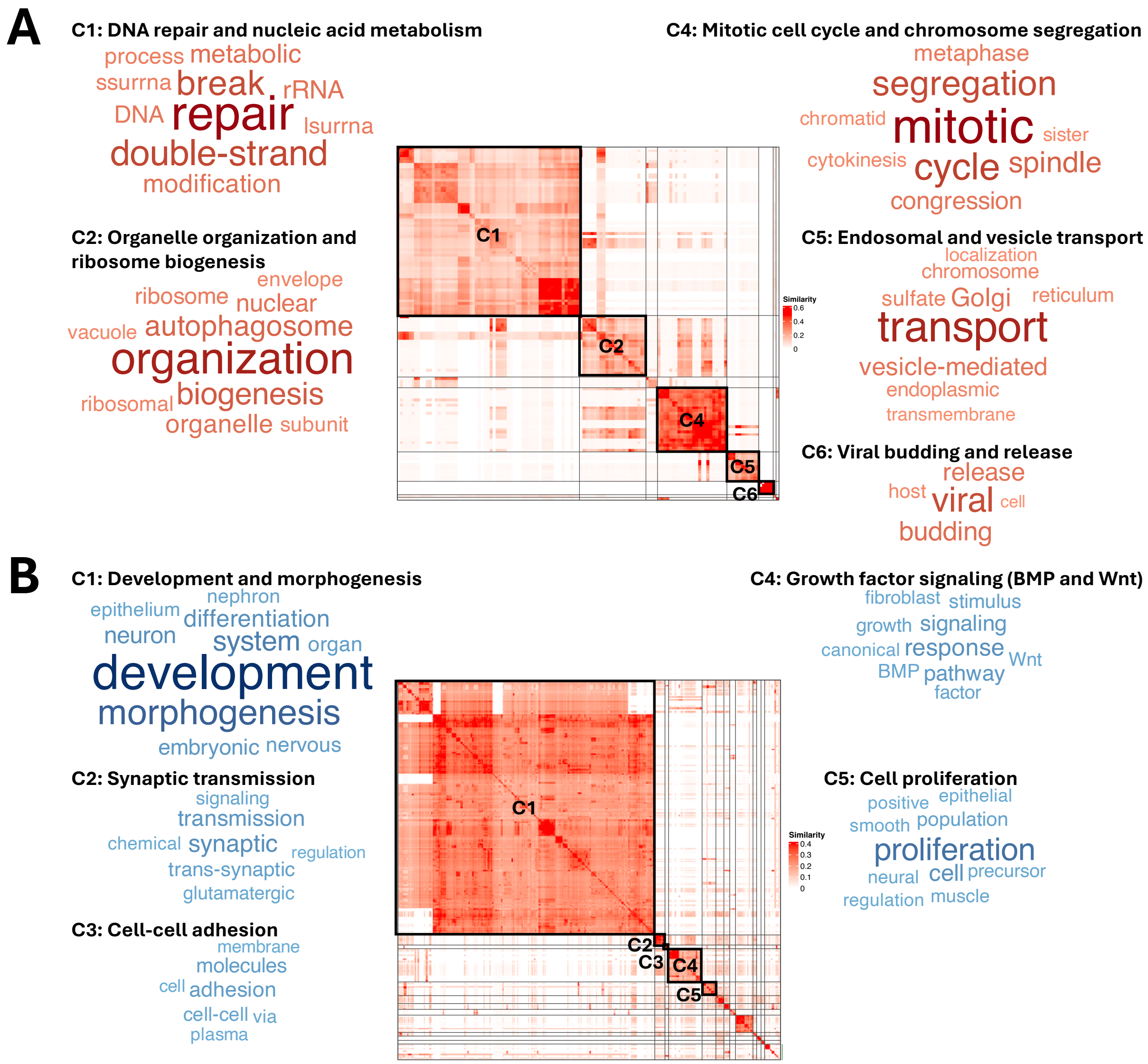


**Supplementary Figure 5. Clusters of GO biological-process terms enriched or depleted in blood BrESS genes.**

(A) Terms enriched in blood BrESS genes (132 terms, FDR < 0.05, NES > 0). (B) Terms depleted in blood BrESS genes (526 terms, FDR < 0.05, NES < 0). Each heatmap shows pairwise semantic similarity among the terms, which were clustered by binary cut. Clusters are numbered in order of the P value of their most significant term. Boxes mark the clusters shown with word clouds (C1, C2 and C4–C6 in A, and C1–C5 in B). Word clouds show the keywords enriched in the term names of each cluster, with up to ten words per cluster, and word size and color intensity indicate $-\log_{10}$ adjusted P. Cluster names are those used in Fig. 5D and Supplementary Table 11. All clusters with at least five terms (5 in A and 12 in B) are shown in Fig. 5D and listed with their member terms in Supplementary Tables 11 and 12.

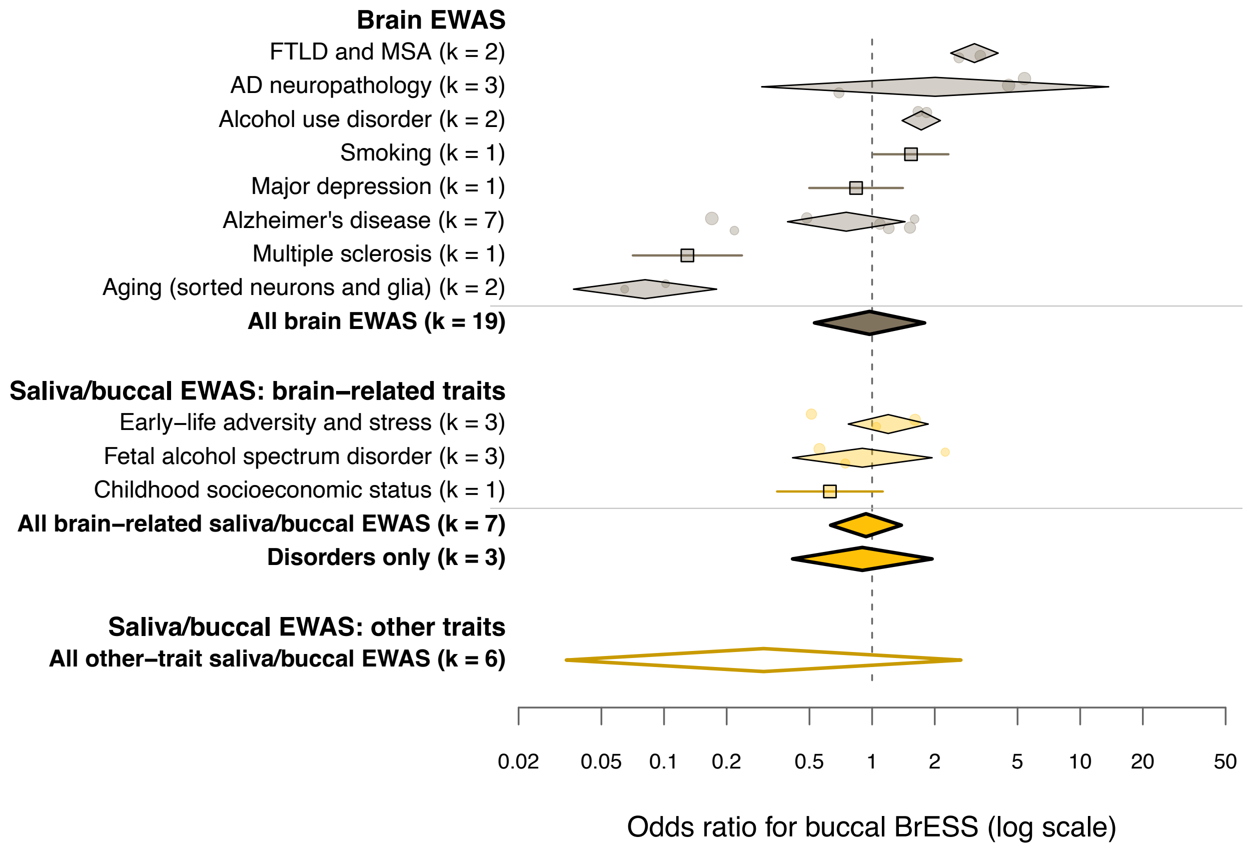


**Supplementary Figure 6. Overlap between published EWAS findings and buccal BrESS across trait groups.** CpGs reported by EWAS in the EWAS Atlas and the EWAS Catalog were evaluated for overlap with buccal BrESS. Studies were grouped by trait within each tissue. Sections show brain EWAS, saliva/buccal EWAS of brain-related traits and saliva/buccal EWAS of other traits. Odds ratios are adjusted for CpG location (relation to CpG island × gene region). Small circles indicate individual studies, and their size reflects sample size on a logarithmic scale. Diamonds indicate pooled estimates for each trait group with 95% confidence intervals. Squares with horizontal lines indicate trait groups represented by a single study. Bold diamonds indicate pooled estimates across all studies in each section. The diamond labelled ‘Disorders only’ indicates the pooled estimate for studies of psychiatric and neurological disorders among the brain-related traits. The open diamond indicates the pooled estimate for saliva/buccal EWAS of other traits. k denotes the number of studies. Colors indicate the tissue of the EWAS (gray, brain; yellow, saliva/buccal). The vertical dashed line marks an odds ratio of 1. The x-axis is on a log scale. Pooled estimates, group comparisons and the list of studies are in Supplementary Tables 17, 18 and 19.

### Supplementary Note 1. Expansion and characterization of AMAZE-CpG2 (DB5)

#### Overview

AMAZE-CpG2 (DB5) is a dataset of paired brain–peripheral DNAm profiles from living human brain tissue and peripheral tissues (blood, buccal epithelium and saliva). It expands AMAZE-CpG (n = 19), reported by Nishitani et al.[18], to n = 35 in the present study by adding 16 newly collected subjects. AMAZE-CpG2 is an expansion and update of the first brain–peripheral DNAm correlation database based on an Asian population, and as such has standalone value as an independent resource. Within the present study, it is used mainly as an independent external-validation cohort for TRACE-CpG (Fig. 4D), and it is also one of the cohorts of the COMPASS meta-analysis (DB5). This note describes the composition and general characteristics of the expanded dataset. The data are also available as a web resource (https://epigen.shinozakilab.net/amaze2.html).

#### Participants and sample collection

Characteristics of all 35 participants are provided in Supplementary Table 5, and the location of each resected brain tissue is shown in Supplementary Fig. 2. Of the 35 participants, the first 19 are reproduced from Nishitani et al.[18], and the remaining 16 are newly collected and reported in the present study. Sample collection and DNA extraction followed the procedures described in the previous study[18].

#### Data processing

All AMAZE-CpG2 samples were profiled on the EPIC v1 array. Preprocessing followed the same procedure used for the EPIC v1 data of the other cohorts in this study (Methods). Based on the PGC PTSD EWAS pipeline, background and dye-bias correction was performed with the Noob method, and probes were removed on the basis of detection P value and missingness, together with cross-reactive probes. No batch correction was applied. AMAZE-CpG2 and the IMAGE-CpG2 dataset were therefore processed on the same basis, which allows them to be compared.

#### Genome-wide characterization

An MDS plot of genome-wide DNAm profiles is shown in Supplementary Fig. 3A (n = 35 subjects with all four tissues, 140 samples in total). Brain samples formed a cluster separate from the peripheral tissues, and the peripheral tissues were arranged in a manner consistent with their cellular compositions. This pattern is consistent with that observed in the previous study and in IMAGE-CpG2.

#### Population stratification

To examine the difference in population background between AMAZE-CpG2 and IMAGE-CpG2, we performed a DNAm-based population stratification analysis (Supplementary Fig. 3B). This analysis was restricted to EPIC v1 samples, comprising AMAZE-CpG2 (35 subjects, all profiled on EPIC v1) and the EPIC v1 subset of IMAGE-CpG2 with blood DNAm (86 subjects). EPIC v2 was redesigned with substantial removal of the SNP-overlapping probes used for ancestry inference by the two methods below, so restricting both datasets to EPIC v1 ensured probe compatibility. Both methods capture population structure by principal component analysis of CpG sites near SNPs. One is the approach of Barfield et al.[S1], the same as in the previous AMAZE-CpG study (5,825 SNP-CpGs). The other is EpiAnceR+ (Höffler et al.[S2]), which residualizes the SNP-overlapping CpG sites for technical and biological factors and additionally integrates the genotype calls from the SNP (rs) probes on the array (3,596 SNP-CpGs). With both methods, AMAZE-CpG2 (Japanese) and IMAGE-CpG2 v1 (predominantly European) were clearly separated.

**Brain–peripheral correlation**

Brain–peripheral DNAm correlations in AMAZE-CpG2 are summarized in Supplementary Table 6 and Supplementary Fig. 3C. Per-CpG Spearman correlations (ρ) between brain and peripheral DNAm were computed across the 35 paired subjects, over 820,406 CpG sites. For blood, buccal epithelium and saliva, we report the mean and median ρ, standard deviation, interquartile range and the proportion of CpG sites with ρ above or below each threshold (>0.2, >0.5, >0.8, <0). To assess whether these correlations are consistent across independent datasets, we compared per-CpG ρ between the IMAGE-CpG2 dataset and AMAZE-CpG2 (Supplementary Fig. 3D). Across 687,798 CpG sites common to both datasets, the two sets of ρ corresponded well (Pearson r 0.62 for blood, 0.60 for buccal epithelium and 0.46 for saliva).

### Supplementary Note 2. Predicted versus measured brain DNAm at representative BrESS

#### Rationale

TRACE-CpG predicts brain DNAm (pBrain) from peripheral DNAm at each BrESS with the model fitted to the full discovery data, using peripheral M-value, age and sex as predictors. Because pBrain is a linear function of the peripheral signal, it adds no information over the peripheral data in group-level comparisons. The predicted brain level itself can nonetheless be of interest, for example to see where peripheral DNAm at a given CpG site sits relative to measured brain DNAm. Supplementary Fig. 4 illustrates this for representative CpG sites, showing on the beta-value scale how the prediction moves blood DNAm toward the range of measured brain DNAm. The figure is a visualization and does not test prediction accuracy, group differences or causality.

#### Selection of the 22 CpG sites

To choose CpG sites of biological interest rather than an arbitrary subset, we intersected the 1,048 CpG sites associated with schizophrenia in the blood DNAm meta-analysis of Hannon et al.[38] with blood BrESS. Twenty-two CpG sites were shared, and all 22 are on the 450K array used for the measured brain dataset. The CpG sites were thus selected for their association with schizophrenia in blood and their BrESS status, not for case–control differences in brain.

#### Variance correction of pBrain

Because pBrain is a linear prediction based on the brain–peripheral relationship, its inter-individual variance is systematically smaller than that of measured brain DNAm. To restore brain-level variance, the deviation of each predicted value from the per-CpG mean of pBrain was multiplied by a scaling factor s, which enlarges the variance while leaving the mean unchanged:

$$\mathrm{pBrain_c} = \mu + (\mathrm{pBrain} - \mu)s$$

where μ is the mean of pBrain across individuals at that CpG. The scaling factor was defined from the discovery data as

$$s = \sqrt{\frac{\sigma_{brain}^2}{\sigma_{pBrain}^2}}$$

where $\sigma^2$brain is the inter-individual variance of measured brain DNAm and $\sigma^2$pBrain is the variance of pBrain, both computed in IMAGE-CpG2. Because s is derived solely from the discovery data, the correction requires no brain measurements from the user. Values of s were precomputed per CpG for each peripheral tissue and are given in the Scale column of Supplementary Data 2. In AMAZE-CpG2, this fixed-variance correction was compared with no correction and with a correction based on the variance of the user's own predicted values (Supplementary Table 7). Without correction, the variance of pBrain was only 32–39% of that of measured brain DNAm, whereas scaling by the user's own predicted variance overshot it by roughly two-fold. The fixed-variance correction brought the ratio closest to unity in both tissues (median 0.73–0.76) with little increase in root-mean-square error, and was therefore adopted. Because the correction rescales deviations around the per-CpG mean, it does not change the ordering of donors, and the per-CpG correlation with measured brain DNAm is the same for all methods.

**Prediction of pBrain**

For each of the 22 CpG sites, pBrain was predicted from blood DNAm (GSE152026, n = 934) with the blood TRACE-CpG model, and the fixed-variance correction derived from the discovery data was applied (see above; Supplementary Table 7). Where chronological age was missing, it was imputed with DNAm-predicted age from the Horvath clock[S3]. The predicted values were converted back to beta values for display. Measured brain DNAm was taken from GSE74193 (Jaffe et al.[4]). Samples were restricted to those that passed the quality control of the original study, had a recorded sex matching the predicted sex, were older than 16 years, were not on plate Lieber_244 and were diagnosed as control or schizophrenia (n = 244). Measured brain DNAm was not used in the prediction.

**Figure elements**

In Supplementary Fig. 4, each panel corresponds to one CpG site and shows three distributions of beta values: measured blood DNAm (Blood), predicted brain DNAm (pBrain) and measured brain DNAm (Brain). Every sample is plotted as an individual point. The overlaid box marks the median and interquartile range (IQR), with whiskers extending to 1.5 × IQR. The y-axis range is set separately for each panel so that the relative positions of the three distributions are visible at each CpG site.

**Interpretation and caveats**

Blood and pBrain (GSE152026, EPIC array) and measured brain (GSE74193, 450K array) come from different cohorts and array platforms. Systematic, non-biological offsets between the predicted and measured brain distributions are therefore expected, and exact numerical agreement is not expected. The shift of pBrain toward the measured brain range reflects the deterministic prediction and should not be read as evidence of a causal relationship. The figure is intended to illustrate, at representative CpG sites, that the prediction moves peripheral DNAm toward measured brain-like values.

**Supplementary Data 1. Leave-one-dataset-out cross-validation results for all CpG sites.**
Each row is one CpG site (*n* = 687,873). For blood and buccal epithelium (columns prefixed Blood_ and Buccal_), R2gain_DB8, R2gain_DB7 and R2gain_DB6 give the gain in out-of-sample $R^2$ for brain DNAm from adding peripheral DNAm to a model with age and sex, in the cycle tested on the named dataset (trained on the other two). n_cycles_R2gain_gt0.2 is the number of cycles with $R^2$gain > 0.2. BrESS is TRUE for CpG sites with $R^2$gain > 0.2 in at least two cycles. Common_BrESS is TRUE for CpG sites that are BrESS in both tissues.

**Supplementary Data 2. TRACE-CpG prediction models for BrESS.**
Linear mixed-model parameters for BrESS in blood and buccal epithelium, estimated on the full IMAGE-CpG2 data (DB6–DB8). Each row is one CpG–tissue combination. R2_Marginal and R2_Conditional are the marginal and conditional $R^2$ of the full model, and R2m_Reduced is the marginal $R^2$ of the reduced model with age and sex only. Coef_Intercept, Coef_Peripheral_M, Coef_Age and Coef_Sex are the fixed-effect coefficients on the M-value scale (sex coded as male = 1). Var_Donor, Var_Region and Var_Residual are the random-effect variance components for donor and brain region and the residual variance. Scale is the factor used to restore brain-level variance in predicted values, computed from the discovery data.
